# Scan field rotation with a K-mirror for fast two-photon imaging along curved planes

**DOI:** 10.64898/2026.10.09.757907

**Authors:** Shinjini Ghosh, Luca Godenzini, Cécile Telliez, Nikita Vladimirov, Lukas Lüchinger, Yuanzhao Liu, Fritjof Helmchen, Peter Rupprecht

## Abstract

In contrast to the curved geometry of biological structures, the imaging plane of conventional microscopes is flat. In principle, axial scanning can adjust the focus to follow curved structures. In resonant scanning microscopes, which are broadly used for video-rate functional imaging, however, such focus adjustment is typically possible at rates faster than the slow (∼30 Hz) scan axis but not as fast as the fast (∼8000 Hz) resonant axis. The fixed direction of the fast resonant axis therefore constrains how the imaging plane can be tilted or curved using axial scanning, and limits how well it can be aligned with biological structures of interest. Here, we present an optical modification that overcomes this constraint. Specifically, we combine scan field rotation using a K-mirror with fast axial scanning using an electrically tunable lens. We demonstrate the versatility of our approach for calcium imaging by aligning the imaging plane with the curved pyramidal cell layer in mouse hippocampus, increasing the number of simultaneously imaged neurons. In addition, we image dendritic arbors of hippocampal and cortical pyramidal neurons that would otherwise be only partially accessible for fast imaging, and we apply our method to the curved layers of cerebellar Purkinje cells. Furthermore, we demonstrate how scan field rotation can be used to optimally orient a dendritic branch for high-speed subcellular glutamate imaging. Our approach is based on a single module and can therefore be retrofitted to standard two-photon systems. We openly share design specifications, 3D-printed parts, and custom ScanImage-compatible code to facilitate rapid adoption.

## Introduction

With an ideal microscope, it would be possible to freely adapt the imaging plane to the structure of interest. In biology and neuroscience in particular, many such structures of interest are not flat and therefore difficult to capture with the flat imaging plane typical for standard microscopes. For example, in mouse hippocampus, the pyramidal cell layer in CA1 and even more so in CA3 forms a curved sheet^1–4^. When such a structure is imaged with a flat imaging plane, only a small subset of cells lies within the focus (Figure 1A), limiting the useful field of view (FOV) and the number of neurons that can be monitored simultaneously. Similar challenges arise in other layered tissues across biology, including cortical or cerebellar layers, the retina, epithelial sheets, or, more generally, for any experiment where efficient sampling depends on aligning the imaging plane with the biological structure of interest (Figure 1A). This alignment, however, is only possible if the scan path can be flexibly adjusted in 3D.

**Figure 1.**
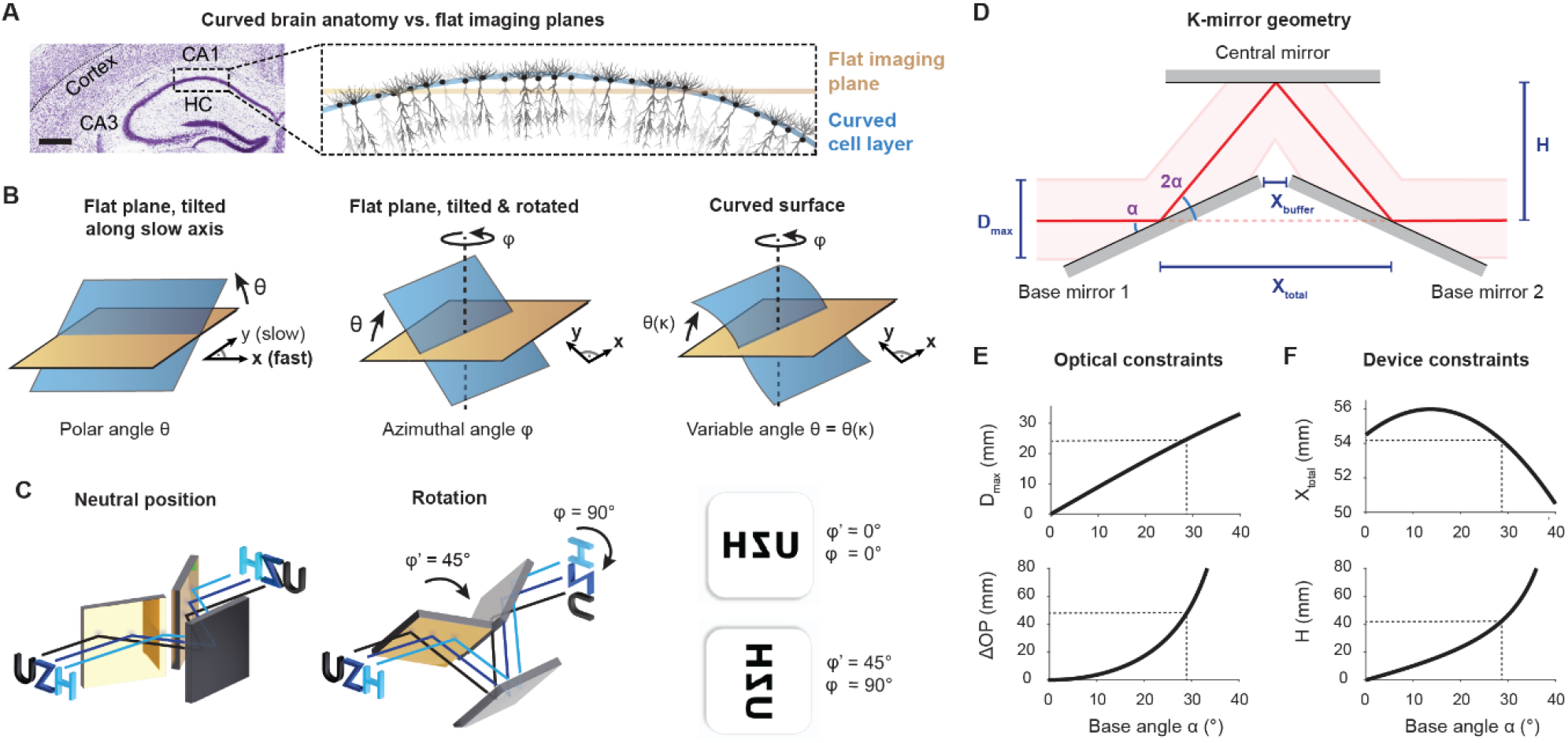
A rotating K-mirror to optimize scan field orientation. **A**, Structures in the brain like cortex or hippocampal CA1 of curved geometries are not well-fitted by a flat imaging plane. The coronal slice image was adapted from the Allen Mouse Brain Atlas^64^. **B**, Rotation of the scan field recovers the degree of freedom missing for resonant-scanning microscopes. *Left:* For resonant-scanning microscopes, the axial change of the image plane (tilt with polar angle θ) is limited to the slow (y) axis. *Middle:* Scan field rotation (azimuthal angle φ) enables tilting along arbitrary directions. *Right:* Parameterization of the local curvature (θ as a function of a parameter κ) to align with locally curved structures. **C**, Scheme of the working principles of a K-mirror. *Left:* An unrotated K-mirror acts as a image-flipping device. *Middle:* Rotation of the K-mirror around the optical axis by φ’ rotates the image by twice the rotation angle φ = 2φ’. *Right:* K-mirror rotation by 45° results in image rotation of 90°. **D**, Geometric and optical parameters of a K-mirror, with base angle *α*, maximal beam diameter *D*_*max*_, distance between mirrors *H*, and the device-specific distances *X*_*total*_ and *X*_*buffer*_. **E**, The base angle *α* determines the effective maximal beam diameter *D*_*max*_ (upper plot) and the additional optical path length *ΔOP*. **F**, *α* determines device dimensions along the optical axis (*X*_*total*_) and along the orthogonal axis (*H*).

In neuroscience, raster-scanning two-photon microscopes are the workhorse for high-resolution *in vivo* imaging^5,6^. These microscopy systems typically rely on a pair of orthogonal scan mirrors – one for fast (x) and one for slow (y) scanning – to produce sequential 2D frames^7^. Earlier designs consisted of two position-controlled galvanometric scan mirrors, with a maximal line rate of ∼2 kHz, resulting in frame rates of ∼4 Hz for 512×512 images. Galvo-galvo systems can follow arbitrary linear combinations of x- and y-scan signals, allowing the raster orientation to be electronically rotated. However, the relatively low line rates limit the frame rates necessary to capture fast neuronal dynamics. During the last two decades, the wide adoption of resonant scanners (line rates of typically 16 kHz) for the fast scanning axis has increased the typical frame rate to ∼30 Hz, which is essential for monitoring of fast physiological processes and neuronal activity across large populations^8,9^. However, this gain in speed comes at the cost of scan flexibility. The scan path in resonant-galvo systems is highly constrained because the x- and y-axes linked to the fast (resonant) and the slow (galvo) scanner are fundamentally different and thus fixed. Consequently, unlike in a galvo-galvo system, the two scan signals cannot be linearly combined to rotate the raster. This constraint poses a fundamental limitation on resonant-scanning microscopes when imaging biological structures.

Z-scanning methods can be used to vary the imaging depth and thereby adapt to biological structures in 3D^5^. Relatively fast axial movement of the focal plane is enabled by a diversity of approaches including objective-attached piezos^10^, electrically tunable lenses^11^ or remote focusing devices^12–16^, with settling times of few milliseconds. Only few highly specialized microscopes based on acousto-optic deflectors (AODs) or linear spatial light modulators (SLMs) are capable of going below this millisecond limit^17–19^, and these implementations are technically more complex and not commonly available on standard resonant-scanning two-photon microscopes. The more typically used axial scanning methods can therefore adjust the axial focus only along the slow scan axis (galvo scanner) but not along the fast axis (resonant scanner). This constraint prevents the experimenter from freely orienting the imaging plane in 3D (Figure 1B, left). Potential solutions to this constraint for resonant scanning systems include physically rotating the scan head, rotating the microscope, or rotating the sample to align the scan plane with the desired biological structure. However, these approaches are mechanically challenging and impractical for routine imaging in vivo.

Here, we describe a simple solution that optically rotates the scan field while leaving both the microscope and the preparation unchanged and thereby extends the flexibility of existing resonant-galvo-based microscopes (Figure 1B). Our approach is based on a rotatable K-mirror optical assembly placed in the beam path. A K-mirror is a device that allows rotation of the entire image without moving any part of the original microscope or the observed object^20,21^. K-mirrors have been used in astronomy in the 20th century as an alternative to Dove or Pechan prisms to cancel the image rotation when tracking stars^22^. Since then, K-mirrors have also been successfully used for microscopes, e.g., for structured illumination microscopy^23,24^. Here, we combined K-mirrors with fast z-scanning – implemented via an electrically tunable lens –, enabling flexible alignment of the microscopy imaging plane with tilted or curved biological structures.

We demonstrate the utility of this method with a diverse set of use cases where structures that would otherwise be inaccessible become accessible for fast imaging. Additional benefits of our approach are that it is easy to implement and compatible with existing microscopes and image acquisition software (e.g., ScanImage), making it a practical and versatile add-on to modern point-scanning systems.

## Results

### Design principles of a K-mirror image rotator

An “image rotator” adds a degree of freedom (rotation) and therefore enables more flexible choice of the imaging plane. One way to build such an image rotator uses a K-mirror that is itself rotated within a mount fixed in the microscope’s scanning beam path. A K-mirror is an optical configuration consisting of 3 flat mirrors that are arranged to form the shape of a “K” with their surfaces^20,21^. The incoming image is flipped by the K-mirror axis. Therefore, a rotation of the K-mirror rotates the flipping axis by a specific angle *φ*′ and thus rotates the image by *φ* = 2*φ*′ (Figure 1C).

The design of a K-mirror must consider two main constraints. First, the K-mirror can only transmit a beam of maximal diameter *D*_*max*_. Second, insertion of the K-mirror assembly adds an additional optical path length, *ΔOP*, and requires physical space, characterized by *X*_*total*_ and the height of the K-mirror *H* (Figure 1D). These values are determined primarily by the angle *α* of the K-mirror’s legs and the size of the individual mirrors used within the K-mirror. When we assume the use of 3 identical square mirrors with side length *L* and thickness *T* (L = 50.8 mm, T = 6.0 mm for 2” Thorlabs mirrors), and a spacing between the two base mirrors of *X*_*buffer*_ (a buffer distance, which we set to 3 mm), we obtain expressions for *D*_*max*_, *X*_*total*_ and *H*:

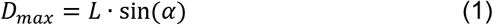

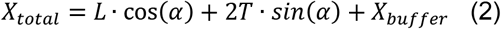

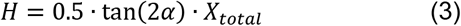

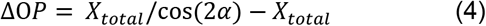

Based on these relationships (Figure 1E,F), we identified a configuration using an angle of *α* = 29° that provides a large effective aperture of the unscanned beam (∼25 mm) to fill high-NA, low-magnification objectives, while increasing the total beam path by *ΔOP* = 47 mm. For applications with different constraints on beam diameter or mechanical geometry of the optical path, the appropriate solution can be based on these equations.

### Implementation variants of a K-mirror image rotator

K-mirror devices have been used before in other microscope types for scan field rotation or to flip the scan field (e.g., in the commercial confocal microscope SP8 by Leica, or refs. ^23,24^). We tested two K-mirror variants that were available to us. First, we used a K-mirror assembly with an effective aperture of 22 mm that was part of a phased-out commercial confocal microscope (Leica SP8), hence called ‘*Leica K-mirror*’ (Figure 2A). Second, we designed a custom K-mirror based on a 3D-printed holder together with off-the-shelf 2” mirrors (Thorlabs) that are slid in and held in place by low-stress silicone glue (Figure 2B). This ‘*3D-printed K-mirror*’ provides a simple-to-test prototype for first users and a potential starting point for future improvement or adaptation. Overall, the Leica K-mirror and the 3D-printed K-mirror exhibited similar characteristics (*α* = 29°; D = 22 mm; Figure 2A,B).

**Figure 2.**
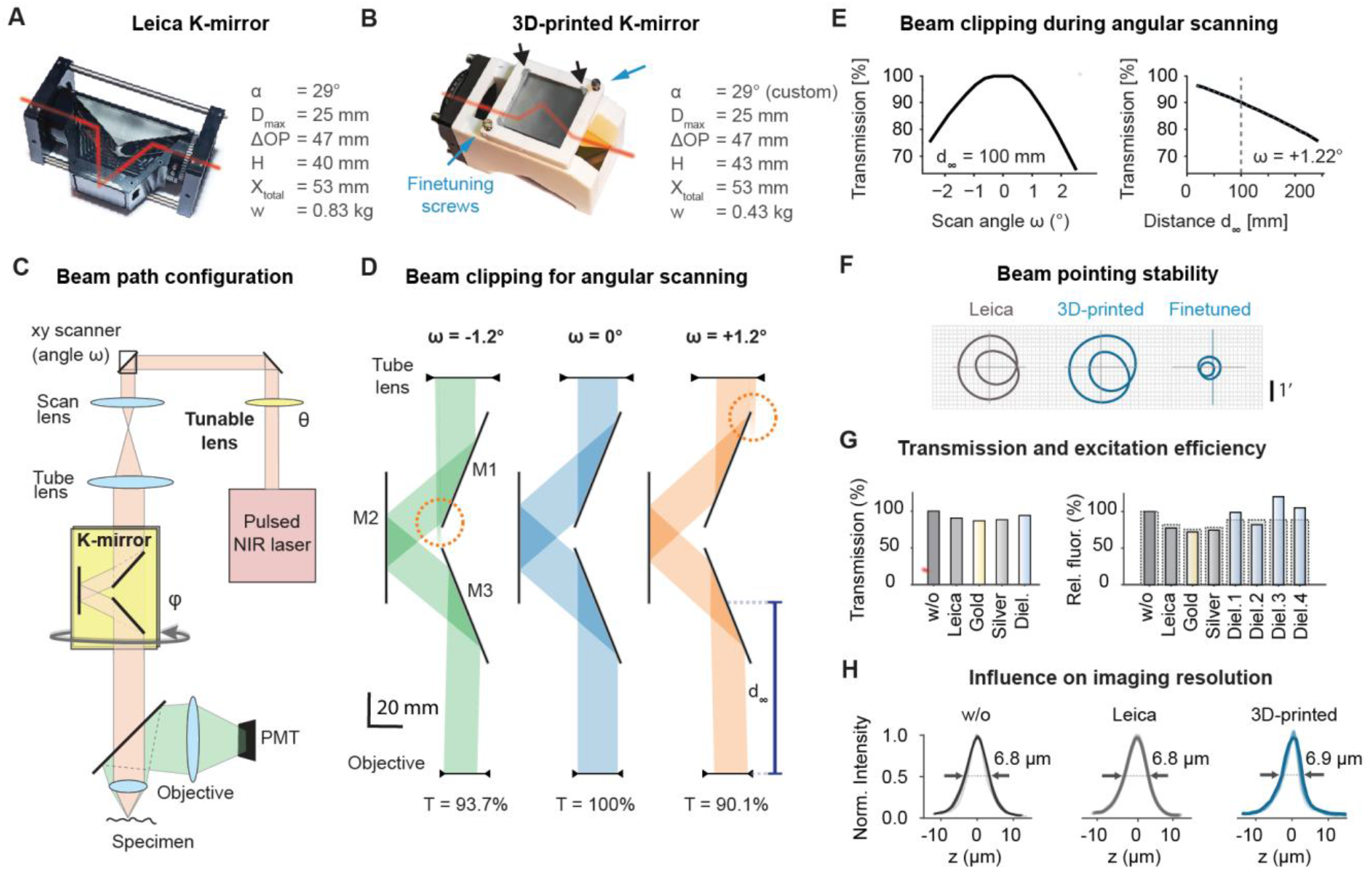
Implementation of a K-mirror image rotator for two-photon microscopy. **A**, K-mirror design of a legacy confocal microscope (Leica SP8). Angle *α*, height *H*, length *X*_*total*_, and weight *w* were measured, maximum acceptable beam diameter *D*_*max*_ and *ΔOP* were computed based on the relationships in Figure 1E. **B**, 3D-printed K-mirror, using square Thorlabs gold mirrors with ∼51 mm side length. Mirrors are slid into the 3D-printed holders and held in place by silicone glue (black arrows). The top mirror can be adjusted via screws for finetuning (blue arrows; cf. Panel (F)). **C**, Microscope configuration for insertion of the K-mirror in the collimated beam path between tube lens and objective (“infinity-space configuration”). The K-mirror controls the azimuthal angle *φ*, an electrically tunable lens controls the polar angle θ. **D**, Scanned laser beam between tube lens and objective at various scan angles (ω, angle taken between tube lens and objective). The beam is only slightly clipped for the maximum scan angles (red dotted circles). **E**, Quantification of beam clipping shown in (D). The quantification assumes a 22-mm diameter top-hat beam. Transmission decreases substantially for large scan angles only (*left*) and can be increased by minimizing the distance d_∞_ between K-mirror and objective. The physical dimensions of the 3D-printed K-mirror result in a minimal d_∞_ of 17.5 mm. d_∞_ = 100 mm represents a practical compromise to reduce beam clipping while leaving space for the dichroic and the axially motorized objective. **F**, Beam pointing stability for the two K-mirrors and with a finetuned 3D-printed K-mirror. For the used microscope, an angle deviation of γ = 1’ corresponds to a lateral translation of 5.7 μm. **G**, *Left:* Transmission of a small laser beam at 905 nm with and without K-mirrors, normalized to without K-mirror. *Right:* Fluorescence yield, relative to without K-mirror (“w/o”), measured using a homogeneous fluorescent plastic slide. Dashed boxes: predictions from transmitted power. Multiple measurements with dielectric K-mirrors resulted from rotation of the K-mirror by 45° increments. **H**, Typical point spread functions obtained for different configurations, with slightly underfilled back aperture of a 10x 0.6 NA objective, measured with 1 μm-beads.

The most straightforward application of such an image rotator is to insert it into the beam path between tube lens and objective of a two-photon microscope (Figure 2C; “infinity-space configuration”). This infinity-space configuration is easy to implement since the laser beam is collimated between tube lens and objective and therefore not sensitive to small changes of the optical distance. Using 3D-printed adapters, the K-mirrors were mounted onto a commercial rotational mount that can be attached to a 2” lens tube. An alternative “scanning-space configuration” is discussed below.

### Characterization of the effects of the K-mirror in the microscope

Despite the simplicity of K-mirrors, it is worth highlighting the practical difficulties that come with their use for two-photon microscopy. First, we used geometric optics to simulate beam clipping for the infinity-space configuration. During scanning, the beam can extend beyond the apertures of the K-mirror, resulting in lower effective transmission. In our case, transmission was reduced only marginally by the K-mirror (>90% transmission for a top-hat beam profile across the scan range: 2.4° peak-to-peak before the objective; Figure 2D,E). In addition, we observed that clipping can be minimized by positioning the K-mirror as close as possible to the objective (Figure 2E) because the beam is clipped primarily by the first mirror of the K-mirror (Figure S1). Since these values may differ largely depending on scan angle range, scan/tube lens and laser beam diameter, we provide a Python notebook that enables users and designers to explore the effects of these parameters on beam clipping (Methods).

Next, we experimentally measured the pointing stability of the Leica K-mirror and the 3D-printed K-mirror (Methods). With careful alignment of either K-mirror variant within an optical cage system, we found that the largest diameter of the curve was approximately 4 arc minutes (4’, 0.067°) (Figure 2F). This angle corresponds to a distance of approximately 21 μm in the focal plane when assuming a 10x Olympus objective (f = 18 mm) and is approximately 36x smaller than the maximal scan angle range for this specific microscope design (2.4° peak-to-peak). Therefore, the alignment errors have a relatively small effect on FOV position.

To theoretically understand pointing stability, we noticed from the analysis of rotational prisms^25^ that the pointing trajectory during rotation can be described by a limaçon of Pascal, with the radial deviation *r* described as a function of the K-mirror rotation angle *φ*′:

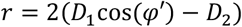

*D*_1_ can be interpreted as the misalignment between mechanical rotation axis and the optical axis of the incoming beam, and *D*_2_ as the misalignment between rotation axis and internal optical axis of the K-mirror assembly. The misalignment errors D_1_, D_2_ are independent from each other and each include both angular and translational components^25^, with the angular component typically dominating. The finetuning screws of the 3D-printed K-mirror, which adjust the tilt of the central mirror, can be used to optimize the *D*_2_ term. In addition to this analytical treatment, we provide a notebook based on geometric optics to interactively explore how the shape of the limaçon is determined by the individual misalignment errors of the K-mirror and by corrective adjustments of the central mirror (Methods). Experimentally, optimizing the finetuning screws of the central mirror reduced the limaçon to an extent of approximately 1’ (0.017°; 5.2 μm in the focal plane; Figure 2F). Together, this analysis explains the shape of the curve that governs rotations of the scan field and quantifies the pointing stability in our microscope.

Next, we analyzed how the K-mirror’s mirror surfaces themselves affect the passing laser beam. To this end, we compared variants without K-mirror, with the Leica K-mirror, and with the 3D-printed K-mirror. For the 3D-printed K-mirror, we used three different sets of mirrors, consisting of protected gold, protected silver, and dielectric mirrors. We found relatively high overall reflectance at 905 nm for all variants (Figure 2G, left; without K-mirror 100% by definition, Leica 91%, Gold 87%, Silver 88%, Dielectric 94%). In addition, we quantified the two-photon excitation efficiency, measured as the fluorescence of a green fluorescent plastic slide. For two-photon excitation one expects the relative fluorescence values to be proportional to the transmission ratio to the power of 2 (dashed lines in Figure 2G, right). Our measured values were slightly lower compared to the predictions (solid bars in Figure 2G, right; Leica 78% measured vs. 82% from theory; Gold 72% vs. 75%; Silver 74% vs. 78%). This slight reduction is expected and likely due to degradation of the spatial beam profile or to temporal dispersion by the mirrors. Interestingly, the performance of the dielectric mirrors was less predictable, with excitation efficiency sometimes lower and sometimes higher than theoretical predictions (Figure 2G, right). We hypothesize that the dielectric mirrors at unusual incidence angles act as chirping mirrors and shorten or extend the temporal beam profile while leaving the power transmission unchanged. Such a chirping effect through dielectric mirrors has been observed before and has been described as strongly wavelength-dependent^26^. We find indeed such a wavelength-dependence for the dielectric K-mirror variant but not the others (Figure S2). Together, we find that a K-mirror introduces moderate losses of transmission and two-photon excitation efficiency. Based on our experiments, we do not recommend dielectrically coated mirrors despite their high transmission since they may affect the temporal pulse shape of femtosecond lasers depending on the rotation of the K-mirror.

An important factor that determines the performance of any microscope is its spatial resolution. In our microscope design with an underfilled 10x/0.60 objective, we measured the microscope’s point spread function (PSF) with an axial FWHM of 6.8 ± 0.4 μm when using 1 μm-beads (mean ± s.d. across n = 17 beads). We found that this resolution was maintained when using the Leica K-mirror (axial FWHM 6.8 ± 0.1 μm across 21 beads) or when using the 3D-printed K-mirror (axial FWHM 6.9 ± 1.1 μm across 22 beads) (Figure 2H). We note that care must be taken to prevent degradation of the optical resolution by inappropriate fixation of mirrors. Due to their relatively large size, these mirrors are prone to bending and thereby can induce aberrations. In an initial 3D-printed K-mirror prototype, mirrors were held in place by spring-loaded screws. We found that even minor strain on the mirror backs induced substantial astigmatism, which could, however, be mitigated by releasing strain on the mirrors (Figure S3). RTV silicone or other low-stress holding mechanisms are therefore essential to prevent aberrations and the accompanying reduction of optical resolution. Together, these analyses demonstrate that K-mirrors can be used without detrimental effects on the resolution, but care must be taken when fixing large mirrors in place.

In specific existing microscopes, the optical configuration may not allow for a K-mirror to be inserted in the infinity space between tube lens and objective. To address this case, we explored an alternative configuration where the K-mirror is inserted between scan lens and tube lens, using a commercial image rotator chosen for its large aperture (IRMU-FD30, ScienceEdge). We refer to this configuration as “scanning-space configuration”. Since the laser beam between scan and tube lens is not collimated, we used a test system where the distance between these lenses was shortened to compensate for the additional beam path introduced by the K-mirror (Figure S4A,B). Geometric optics simulations showed that beam clipping in our specific microscope configuration is minimal except for scan angles that are beyond the practically used range (Figure S4C-E); we provide a notebook for interactive exploration of beam clipping using the scanning-space configuration for other settings or beam diameters, scan/tube lenses, and K-mirror variants (Methods). Especially for high-NA large-FOV microscopes, which require both large beam diameters and large scan ranges, the scanning-space configuration should be advantageous since the reduced effective beam size in the non-collimated space can be used to reduce beam clipping by choosing an optimized position of the K-mirror along the optical beam path. However, it is also important to note that the scanning-space configuration does not allow for flexible removal or addition of the K-mirror since the physical distance between the lenses has to be set differently for with or without K-mirror.

Together, our measurements and optical simulations demonstrate that high performance in terms of fluorescence excitation and resolution is maintained when either the Leica K-mirror or the 3D-printed K-mirror is introduced but that a few precautions have to be considered. In addition, we provide notebooks on GitHub to explore custom optical configurations and the resulting beam clipping for adaptation and redesign of new microscope configurations (Methods).

### Scan field optimization for imaging curved laminar structures in hippocampus and cortex

To demonstrate the applicability of K-mirror-based scan field rotation paired with ETL-based field curvature, we performed two-photon calcium imaging in hippocampal CA1, targeting the pyramidal cell layer for somatic population imaging. We expressed the genetically encoded calcium indicator GCaMP8m in pyramidal neurons and prepared a cannula-based transcortical window positioned above dorsal CA1 (Figure 3A). The CA1 pyramidal cell layer forms a curved sheet along the transverse axis, with basal dendrites extending into *stratum oriens* (SO) and apical dendrites into *stratum radiatum* (SR) (Figure 3B). As a result of the curvature, a large number of the pyramidal cell bodies can lie partially or fully out of the focal plane, with segments of apical and basal dendrites visible on both sides of a stripe of cell bodies (Figure 3B-C).

**Figure 3.**
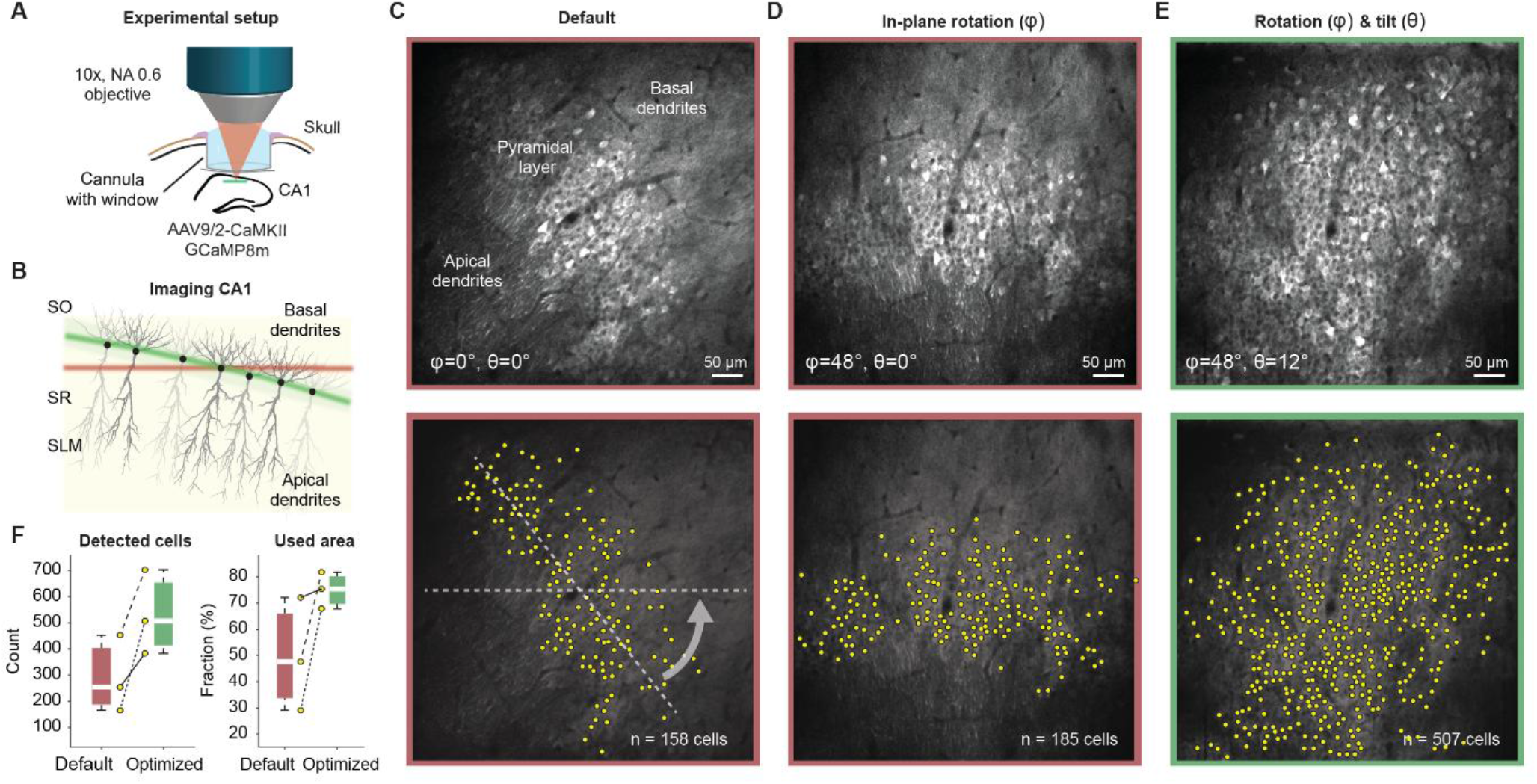
Scan field optimization for imaging of the pyramidal cell layer in hippocampal CA1. **A**, Schematic of experimental setup. A cannula with coverslip is implanted to image excitatory pyramidal cells in hippocampal CA1 of mice in vivo. **B**, Pyramidal neurons in CA1 are arranged in a curved 2D sheet with basal dendrites above and apical dendrites below this pyramidal cell layer. If the curvature of the cell body layer does not align fully with the microscope’s focal plane, only a subsection of the image will contain cell bodies. **C**, Typical FOV for in vivo imaging of pyramidal neurons in CA1. The pyramidal layer appears as a diagonal stripe in the FOV, with individual apical trunks being visible in the lower-left quadrant, and fine blurry basal dendrites visible in the upper-right quadrant (top). Manually detected cell bodies (bottom). **D**, Rotation of the FOV (*φ*) to align the curvature of the pyramidal cell layer with the slow axis (y-axis) of the two-photon scanning microscope. **E**, FOV rotation (*φ*) and image plane curvature (θ) to maximize the number of simultaneously recorded pyramidal cells in an individual imaging plane. F, Quantification of number of visually detected cells (left) and the area of the FOV that contains somata of pyramidal neurons (right) across FOVs recorded with different mice (n = 3 FOVs from 3 mice).

To compensate for this mismatch between the somatic layer and the imaging plane, we rotated the scan field using the K-mirror inserted in the infinity-space configuration (azimuthal angle *φ* = 48°, zenith angle θ = 0°; Figure 3D) to align the curvature of the pyramidal cell layer with the slow y-axis of the laser scanner. To follow the layer’s curvature with axial scanning, we designed a graphical user interface communicating with ScanImage that allows to define a free-form z-scanning trajectory that is only restricted by the temporal responsiveness of the z-scanning device (Methods). For the relatively small FOVs used here (500-600 μm side length; 1024×1024 pixels at 15 Hz), we noticed that the curvature of the hippocampal cell layer can often be well approximated with a sawtooth trajectory for axial scanning, resulting in a simple tilt of the imaging plane along the y-axis (*φ* = 48°, θ = 12°; Figure 3E; Figure S5A,B; Supplementary Movie 1) to align the pyramidal cell layer with the imaging plane. In other cases, more complex scanning trajectories were necessary, for example, to follow a convex curvature of the pyramidal cell layer (Figure S5C). The alignment of scan trajectories with the pyramidal cell layer resulted in a substantial increase in the number of detected neurons (from 291 ± 147 to 531 ± 161, mean ± s.d. across n = 3 animals) and an increase of the effective imaging area containing somata (from 50 ± 22% of full FOV to 75 ± 7%) (Figure 3E,F).

We performed similar experiments also using another microscope designed in a scanning-space configuration (cf. Figure S4), with the K-mirror inserted between the scan and tube lens. To this end, we optimized the scan field to maximize the number of simultaneously imaged pyramidal cells in one animal. Pyramidal cells across the relatively flat part of the CA1 pyramidal cell layer and the curved part of CA2/CA3 were distributed across >150 μm in z (Figure S6A) but were recovered in one imaging plane when optimizing scan field rotation and curvature (Figure S6B,C), resulting in the recording of calcium imaging data from neuronal somata across the CA1 to CA3 subfields of hippocampus (Figure S6D). These results demonstrate the practical utility of the scanning-space K-mirror configuration and show that simultaneous correction of in-plane rotation and curvature can improve the capture of the curved CA1-CA3 pyramidal cell layer.

To showcase the broader applicability of this approach, we applied the same optimization strategy to image in one animal long range axonal projections within cortical layer 1 (L1). In such preparations, the imaging plane of the two-photon microscope can be misaligned with the cortical surface due to the natural curvature of the cortex (Figure S7). Optimization of FOV rotation and curvature (*φ* = 51°, axial tilt by θ ≈ 17°) resulted in improved alignment of the dura as well as axons in L1a and L1b with the scanned imaging plane (Figure S7). This proof-of-concept experiment illustrates that appropriate scan field rotation combined with axial scanning can improve the disentanglement of laminar structures not only in hippocampus but also in the cortex.

### Scan field optimization for dendritic imaging in hippocampal CA1

Functional dendritic imaging aims to capture dendritic activity across a large part of the dendritic tree, together with somatic activity. However, with two-photon raster scanning, it is typically challenging to image a contiguous part of a neuron’s dendritic tree since it is rarely aligned with the scan field. As a result – unless scanning of arbitrary trajectories is enabled by special microscopes^27–29^ – dendritic imaging is constrained to only somata and their surroundings, or to small dendritic fragments imaged across multiple imaging planes^30–34^, unless the dendritic tree happens to be aligned with the imaging plane^33,34^. Here, we demonstrate that adjusting scan field rotation and curvature can help visualize the dendritic structure more completely. To this end, we used the infinity-space configuration and applied K-mirror rotation together with ETL-based z-scanning to visualize GCaMP8m expression in the CA2/3 pyramidal cell layer of a single mouse (Figure 4A-B).

**Figure 4.**
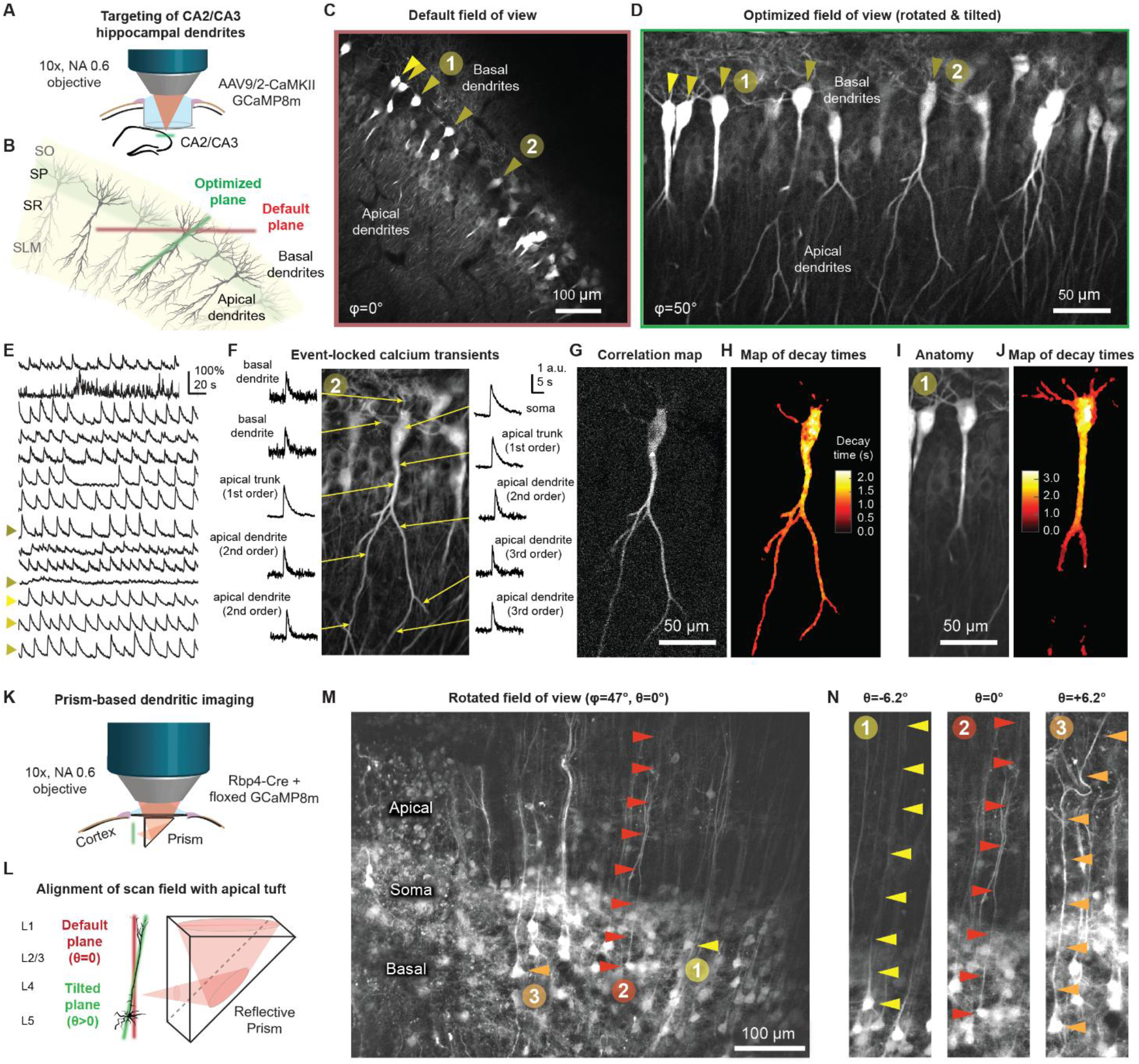
Scan field optimization for dendritic imaging. **A**, Schematic of experimental setup for hippocampal imaging of excitatory pyramidal cells in hippocampal CA2/3 of a mouse in vivo. **B**, The sheet of pyramidal neurons in CA2/3 is arranged in an angle with respect to the imaging plane (red). Tilting the imaging plane along the correct angles *φ* and θ would enable imaging along the dendritic trunk of the neuron (green). **C**, Default imaging plane, with the pyramidal cell layer as a thin stripe, with short fragments of apical dendrites in the lower-left quadrant. **D**, Imaging plane after scan field optimization using scan field rotation (*φ*) and flexible scan plane curvature (defined by variable local tilt angles ranging from θ = 25° at the soma and θ = 0° at distal apical dendrites, covering a total of ∼125 μm in z). Arrow heads indicate neurons identified in (C). **E**, Example neuronal activity of cell bodies of the FOV in (D), recorded during 1.5% isoflurane anesthesia (hence the regular oscillatory activity). Arrow heads indicate neurons highlighted in (D). **F**, Subcellular calcium signals, averaged across soma-detected events (n = 22). Fine apical and basal dendrites exhibit substantially faster calcium transients than somatic or large apical compartments. **G**, Map of pixels that are highly correlated with the neuron’s somatic activity. This correlation map is thresholded and used to define the cellular area for analysis in (H). **H**, Subcellular decay times of an example pyramidal neuron, mapping the half-decay time during somatic calcium events (n = 22) for each pixel. **I**,**J**, Same analysis as in (F-H) for a pyramidal neuron where the basal dendrites were partially included in the imaging plane (based on n = 25 events). **K**, Schematic of experimental setup for cortical dendritic imaging. A coverslip with an attached glass prism is implanted to image L5 pyramidal cells (Rbp4-Cre mice with AAV-based delivery of Cre-dependent GCaMP8s) in the primary somatosensory cortex in vivo. **L**, Apical dendritic trees of L5 pyramidal neurons are arranged approximately in parallel to the surface of the prism, but are not fully aligned with the imaging plane. **M**, After rotation of the FOV, only few dendrites are aligned with the imaging plane across the full extent of the dendrite. **N**, Using the tunable lens to tilt the imaging plane along the y-axis can bring the apical dendrite of specific neurons into the FOV for different tilts (θ = ±6.2°, maximum axial shift of ±60 μm), while keeping the same imaged somata in focus. Arrowheads indicate the soma and apical trunk.

With the default scan field, the pyramidal cell layer appeared as a diagonal strip across the FOV, with apical dendrites truncated in the lower-left and basal dendrites truncated in the upper-right quadrant (Figure 4C), as is often the case for CA3 imaging^1,4,33,35^. By adjusting within-plane axial scanning (maximal local tilt ∼25° around the somata, ∼0° around the distal apical dendrites; full range of ∼125 μm) to match the orientation of the dendritic trunks, we were able to resolve long continuous stretches of both apical and basal dendrites along with the somata across the entire FOV (Figure 4D; 1024×1024 pixels at 15 Hz). Several neurons identifiable in the default configuration (arrowheads, Figure 4C) were recovered with more complete morphology after rotation and applying field curvature (Figure 4D; Supplementary Movie 2).

With calcium data recorded across the dendrites of individual neurons during anesthesia, we extracted transients triggered by somatic events across the subcellular arborization (Figure 4E,F; Methods). Fine apical and basal dendrites exhibited faster calcium transients compared to soma and larger apical trunk compartments, consistent with the known dependence of calcium kinetics on the surface-to-volume ratio of neuronal compartments^36^. To quantify this compartment-specific functional property, we defined a neuronal mask based on a thresholded map of local correlations between somatic transients and pixel-wise fluorescence changes^37^ (Figure 4G). Within this mask, we mapped the event half-decay times on a pixel-wise basis for the visible apical (Figure 4H) or basal (Figure 4I,J) dendrites. We found that half-decay times differed by a factor of up to 5 between soma and dendrites (within-neuron ratio between 5% and 95% percentiles of decay times: 4.0 ± 0.6 across n = 4 neurons). This ratio is similar to ratios of 3-4 observed in hippocampal brain slices with organic calcium dyes^38^. These analyses demonstrate that coordinated K-mirror rotation and ETL-based tilt correction can substantially improve coverage of dendritic imaging and enable spatially precise characterisation of compartment-specific calcium dynamics in hippocampal pyramidal neurons.

### Scan field optimization for prism-based dendritic imaging of pyramidal cells

Prism-based imaging can be used to effectively record from hippocampal or cortical pyramidal cell dendrites that are otherwise orthogonal to the imaging plane^39–41^. However, the imaging plane may still be misaligned with the orientation of the dendritic arbor due to tilt of the prism or due to slightly oblique dendrites. Here, we show that our approach based on FOV rotation and curvature optimization is suitable for addressing this problem. To this end, we performed prism-based imaging of cortical layer 5 pyramidal neurons *in vivo* in one Rbp4-Cre mouse injected with viruses to induce Cre-dependent GCaMP8m expression (Figure 4K,L).

The implanted reflective glass prism was positioned beneath a cranial coverslip and enabled optical access to somata, basal dendrites, and proximal apical dendrites across cortical depth (Figure 4M). Most apical dendritic arbors were arranged nearly parallel to the prism surface. However, local variations in their tuft arborizations caused dendrites to be misaligned with the default imaging plane. We therefore combined scan field rotation with ETL-driven z-scanning along the slow scan axis. Relative to the rotated reference plane, we applied maximal axial offsets of approximately -60 µm or +60 µm across the scan field, while recording from the same somata. With this approach, we were able to selectively bring different contiguous apical trunks into focus at different depths (Figure 4N).

To more broadly confirm the benefit of scan field optimization for prism-based imaging, we inserted a reflective prism into the hippocampus of a mouse and performed calcium imaging of CA1 pyramidal neuron dendrites. As for cortical prism-based imaging, we rotated the imaging plane to align the slow y-axis with the dominant dendritic orientation and showed that it is possible to selectively bring individual apical dendrites into focus at different axial inclinations (Figure S8). Together, these cortical and hippocampal prism experiments demonstrate that our method allows us to optimize the scan field for targeted subcellular imaging of extended dendritic structures.

### Imaging of curved Purkinje cell layers in cerebellar cortex

The mouse cerebellar cortex is a layered and strongly curved brain structure. To demonstrate imaging of curved structures with multiple parallel imaging planes, we performed calcium imaging of Purkinje cells expressing GCaMP8m through a cranial window in one mouse (Figure 5A,B). Because Purkinje-cell somata are arranged along a curved cell body layer, both dendritic structures and Purkinje cell somata were visible across the majority of imaging planes within a z-stack spanning 120 µm (Figure 5C). Thus, no single flat imaging plane captured the entire somatic or dendritic cerebellar layer.

**Figure 5.**
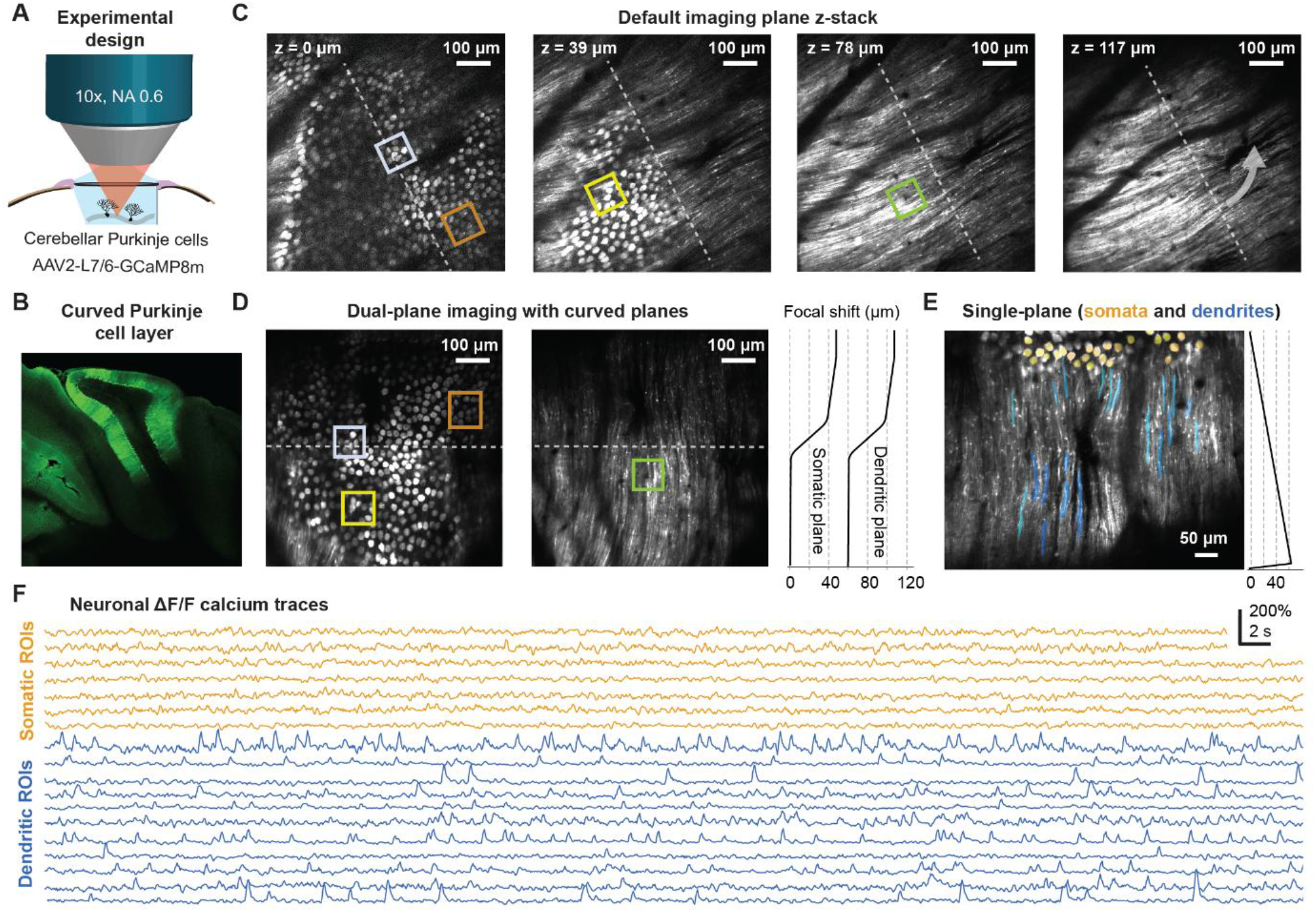
Scan field optimization for the curved layers of cerebellar cortex. **A**, Experimental configuration to image cerebellar Purkinje cells through a cover slip. **B**, Coronal confocal histology image (AP ™5.8 mm from bregma), highlighting the curved Purkinje cell layers across vermis, simple lobule (SIM), and crus1. Green is GCaMP8m fluorescence. Same single animal as in panels (C-F). **C**, Sections of a z-stack from the start of the Purkinje cell layer (0 and 39 μm) to their more superficially located dendrites (78 and 117 μm). Colored boxes highlight cell bodies relocated in panel (D). **D**, Plane-hopping to capture somata (left) and dendrites (right) with curved imaging planes (focal shift indicating the z-scan trajectories) after rotation of the field of view (1024×1024 per frame, volumetric rate of 7.5 Hz). **E**, Single-plane imaging, with the imaging plane rotated to include both somata and their dendrites in the same imaging plane. Z-scan trajectory shown as in panel (D). **F**, Neuronal activity extracted from a subset of ROIs shown in (E). Purkinje cell somata do not show visually detectable calcium transients; dendrites exhibit fast, putative climbing fiber-evoked transients.

We therefore rotated the imaging plane to align the dominant orientation of the cerebellar lobules’ curvature with the slow scan axis and applied axial scan paths to follow the curvature of the somatic layer. Together with plane-hopping, this approach enabled imaging of two distinct curved surfaces corresponding to the dendritic and somatic layers (Figure 5D; two planes of 1024×1024 pixels at 7.5 Hz). This proof-of-concept demonstrates how two parallel curved imaging planes can be used to quasi-simultaneously sample from depth-separated compartments.

To highlight how imaging plane orientations can be chosen with a high degree of freedom, we used the same cerebellar window and adjusted the scan field rotation and z-scanning to record from both somata and dendrites (Figure 5E). This produced a single optimized plane containing multiple Purkinje-cell somata together with their extended dendritic segments. ΔF/F traces from dendritic ROIs exhibited frequent fast transients that are likely to correspond to climbing fiber-evoked dendritic spikes^42–44^ (Figure 5F; 1024×1024 pixels at 15 Hz; Supplementary Movie 3). Somatic calcium signals have been demonstrated to reflect changes in simple spike rates^45^. In our experiments, somatic ROIs did not exhibit visually detectable calcium signals, which may be explained by limited calcium spread between compartments and the higher surface-to-volume ratio of dendrites.

Together, these experiments establish that coordinated scan field rotation and ETL-driven axial scanning can transform a conventional flat raster scan into imaging surfaces that follow the curved three-dimensional organization of cerebellar Purkinje-cell somata and dendrites.

### High-speed glutamate imaging in individual dendrites

Optical sensors that report glutamate concentration changes and membrane potential changes can resolve subthreshold and subcellular activity in neurons but require high-speed imaging and high photon yield to capture fast events^46,47^. With resonant scanning microscopy, high-speed imaging can be achieved using an elongated FOV consisting of few or even a single scan line. However, the fixed orientation of the fast scan axis is typically not aligned with the structure of interest such as dendrites. Here, we demonstrate how scan field rotation using a K-mirror aligns the elongated FOV with the elongated dendritic structure to enable high-speed glutamate imaging.

We expressed the glutamate sensor iGluSnFR4s (ref. ^47^) in layer 5 pyramidal neurons of two Rbp4-Cre mice (Methods) and performed two-photon imaging in posterior parietal cortex (PPC) of anesthetized animals. A cranial window provided optical access to branches of apical dendrites oriented parallel to the cortical surface. However, individual branches were not aligned with the x-axis of the raster scan, requiring large and inefficient FOVs and therefore slow frame rates of ∼30 Hz to fully cover the dendrite (Figure 6A). We therefore rotated the scan field to align the dendritic segment with the fast axis of the FOV. This configuration allowed us to image an extended dendritic segment within a narrow, 48-line imaging strip, which increased the frame acquisition rate to approximately 317 Hz and thereby enabled high-speed sampling of fast glutamate transients (Figure 6B, top).

**Figure 6.**
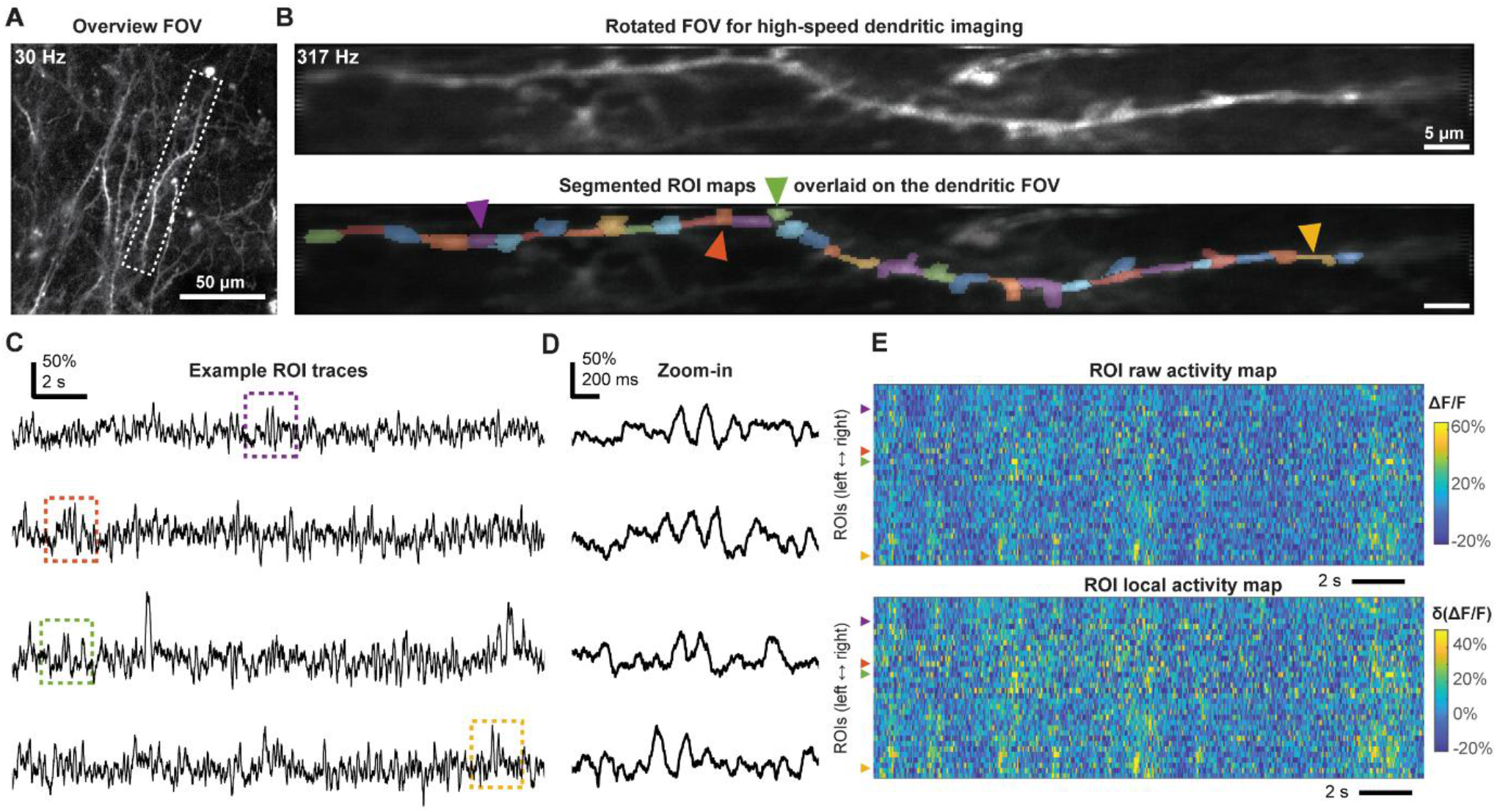
Scan field rotation for high-speed dendritic glutamate imaging. **A**, Imaging of iGluSnFR4s-expressing layer 5 pyramidal neurons in a mouse during anesthesia, overview with slow scanning (30 Hz, 512×512 pixels). **B**, *Top:* Rotated FOV of the dendritic segment highlighted with a dashed rectangle in (A), recorded at high speed (317 Hz, 512×48 pixels), averaged over 30,000 frames. *Bottom:* Colors indicate different ROIs from the dendritic process, covering both dendritic segments and spines. **C**, ΔF/F traces from four example ROIs (marked with arrows in (B)). **D**, Zoom-in to selected local events. **E**, Raster plots of all the ROIs ordered from left to right along the dendritic process in (B). *Top:* Raw ΔF/F of all ROIs. *Bottom:* ΔF/F after subtracting the regressed global fluorescence component.

The optimized FOV was manually segmented into subcellular ROIs along the dendritic shaft (Figure 6B, bottom). iGluSnFR signals exhibited heterogeneous activity patterns, with the majority of events spanning multiple ROIs and others restricted to spatially localized regions (Figure 6C,D). To better isolate spatially confined local events, we subtracted from each ROI the activity shared with the remaining ROIs using linear regression. The resulting residual traces retained spatially restricted, local fluorescence events (Figure 6E). We repeated the same experiments and analyses for a dendrite from another animal (Figure S9). These results demonstrate that scan field rotation enables precise alignment of elongated imaging regions with the elongated geometry of dendritic segments for high-speed imaging of glutamate. This example highlights the practical advantage of K-mirrors to optimize the scan geometry even in the absence of axial scanning.

## Discussion

Here, we introduce the K-mirror as an add-on for existing fast two-photon raster-scanning microscopes. This modification allows to flexibly rotate the field of view and, when combined with a remote z-scanning device (tunable lens), to apply a flexibly curved scan field that matches the biological structure of interest. We demonstrate how this simple approach can optimize the imaging plane, thereby increasing the number of simultaneously imaged cells and enabling more complete or rapid imaging of dendritic structures of individual neurons.

K-mirrors are well established optical devices^20^ that are mainly used in astronomy but also in microscopy. For example, they have been part of commercial microscopes (e.g., the Leica SP8 confocal microscope) and have been adapted for special applications, e.g., to switch between the imaging modes of structured illumination microscopy^23,24^. Here, we make three main contributions. First, we identify the practical value of K-mirrors in enhancing the flexibility of fast resonant-scanning two-photon microscopes. Second, we investigate key aspects of their practical implementation, test in experiments and via optics simulations both commercial and custom-designed K-mirrors as well as different microscope configurations, and provide open source designs for 3D-printing. Third, we demonstrate the utility of K-mirrors for scan field optimization across applications ranging from neuronal population imaging to dendritic imaging to fast glutamate imaging. The breadth of these use cases highlights the broad potential of this technology as an enhancement for two-photon microscopes.

K-mirror-based scan field rotation, even without fast z-scanning, is likely to have immediate applications even beyond the specific use cases demonstrated here. For example, fast voltage imaging with resonant scanners^46^ could be improved by rotating the FOV to maximize the number of neurons. Similarly, line-scan imaging to perform blood flow measurements^48,49^ will benefit from the flexibility to orient the line axis as we have shown for glutamate imaging. In experiments where the head-post of a head-fixed mouse is rotated relative to the microscope body to render navigation more natural^50^, K-mirrors would be the natural solution to de-rotate the FOV online, similar to the original use of K-mirrors in astronomy. Finally, the ability to flexibly follow the curvature of brain structures could be even more important for smaller brains, e.g., of small fish or *Drosophila*, where individual brain regions exhibit strong curvature due to their relatively small size^51–53^.

To facilitate adoption by the community and commercial vendors, we provide CAD files for 3D-printed K-mirrors, which can be directly printed or adapted in CAD programs for specific microscope configurations; we highlight potential practical challenges of using K-mirrors (Figure S2 & S3); and we discuss and test different optical configurations (Figure 2; Figure S4). Furthermore, we share optics simulations of K-mirror microscope configurations as notebooks via GitHub (https://github.com/HelmchenLabSoftware/K-mirror), enabling users to interactively explore these configurations and adapt the design to their own microscope constraints. In general, we found that the infinity-space configuration (Figure 2) provides the simplest integration because the K-mirror can be inserted without modifying the existing beam path. In cases where the space between tube lens and objective is not accessible, the scanning-space configuration (Figure S4) can be advantageous but requires adjustment of the scan or tube lens positions. This configuration may, however, be attractive for high-NA large-FOV microscopes, which require both large scan angles and large laser beam diameters. Since the laser beam size is relatively small around the focal point between scan and tube lens, this may be the natural position of a K-mirror module for high-NA large-FOV microscopes. Finally, to facilitate adoption, we provide code and instructions on GitHub for integrating within-plane axial scanning into the standard microscope control software ScanImage^54^ (https://github.com/HelmchenLabSoftware/K-mirror).

To further simplify the use of K-mirrors, we see several lines of potential further development. First, both custom and commercial microscope design could consider K-mirror insertion already during the design process to facilitate integration, for example by using motorized rotation stages that control and communicate the coordinate system changes with the imaging software. Second, axial focus shifts in our implementation were achieved using electrically tunable lenses. These devices have settling times on the order of a few milliseconds^11^, which can introduce artifacts during rapid focus transitions, especially following large steps. This limitation could be overcome by faster z-scanning technologies with shorter settling times^12–14,16^. Additional limitations exist, which are, however, not specific to our method but are relevant for any technology that results in non-flat imaging planes: Imaging with tilted and curved planes can directly increase the sensitivity to motion artifacts. This is a consequence of the anisotropic optical resolution in two-photon microscopes, with the axial extent of the PSF being larger than the lateral extent. In general, it is possible to post-hoc correct for brain motion artifacts that occur within the imaging plane, but not for brain motion artifacts that occur out of plane. Therefore, if the out-of-plane direction aligns with the axial direction of the PSF, the negative effects of brain motion are only moderate. For curved and tilted planes, however, the out-of-plane direction aligns to a larger extent with the lateral directions of the PSF, resulting in a higher sensitivity to brain motion artifacts. As an illustrative limiting case, a plane tilted by θ = 90° would be highly sensitive to motion, as movements on the order of the lateral resolution (<1 μm) would already result in detectable image shifts. This issue must be considered during experimental design for fast imaging with non-flat planes in a living animal, but it could also be addressed experimentally by isotropic PSF-shaping^55^.

In conclusion, we believe that the simplicity of the K-mirror, together with its practical value across a broad set of applications, makes it ideally suited for rapid adoption by the neuroscience community. Here, we provide all essential analyses, resources and demonstrations to support this integration of K-mirrors as a standard component for two-photon microscopy.

## Acknowledgements

This work was supported by the Swiss National Science Foundation (Ambizione Grant PZ00P3_209114 to P.R.), the University Research Priority Program (URPP) “Adaptive Brain Circuits in Development and Learning” (AdaBD) of the University of Zurich (UZH) (to F.H. and N.V.) an UZH Postdoc Grant (FK-25-057 to C.T.), and the UZH FAN (FAN Fellowship to P.R., supported by the Gebauer Foundation and the Ernst Göhner Foundation). We thank Maria Laura Moreira for help with confocal microscopy, and Philipp Bethge for managing mouse lines.

## Contributions

P.R. conceived and supervised the project. L.L., S.G., and P.R. designed the 3D-printed K-mirror holder. N.V. created the adapter for the Leica K-mirror, performed optical simulations and contributed to optical characterization. P.R. and C.T. constructed microscopes. P.R. adapted ScanImage code. S.G., L.G., C.T., and P.R. performed experiments. S.G., C.T., and P.R. analyzed data. Y.L. contributed to experiments and analyses. F.H. provided resources and guidance for the project. S.G. and P.R. wrote the initial draft of the manuscript. All authors contributed to the writing of the final version of the manuscript.

## Methods

### Animals

All animal procedures were conducted in accordance with the Swiss federal guidelines for the use of animals in research and were approved by the Cantonal Veterinary Office of Zürich. All animals were group-housed under a reversed 12 h light/12 h dark cycle in temperature- and humidity-controlled conditions (21-23°C; 55-60% relative humidity), with food and water available ad libitum.

Experiments were performed using wild-type C57BL/6 mice to target hippocampal pyramidal cells related to Figures 3 and 4, axons in cortical layer 1, and cerebellar Purkinje cells. Rbp4-Cre mice were used to selectively target cortical layer 5 pyramidal neurons, and Thy1-GCaMP6f transgenic mice for hippocampal calcium imaging related to Figure S8. Both male and female mice were used.

### Surgery

#### General anaesthesia, analgesia, and postoperative care

4- to 6-month-old mice were used for experiments. Depending on the experimental protocol, analgesia was provided with either Metacam (5 mg/kg body weight, s.c.) and Buprenorphine (0.1 mg/kg, s.c.) (protocol 1) or buprenorphine extended release (Ethiqa XR, 2.5 μl/g body weight, s.c.) (protocol 2) before surgery. Anaesthesia was induced using isoflurane (5% in O_2_ for induction, 1-2% for maintenance during surgery), and the body temperature was maintained at 35-37°C using a heating pad. Ophthalmic ointment was applied to prevent corneal drying.

After surgery, animals were monitored postoperatively. Depending on the analgesic regimen, mice received either buprenorphine in the drinking water for 72 h with scheduled subcutaneous buprenorphine injections (protocol 1) or continued release from Ethiqa XR (protocol 2). Calcium imaging was performed 2-5 weeks after the surgery.

#### General surgical preparation

For all surgeries, shaving cream was applied to the dorsal head above the brain, and an incision was made into the skin after local application of lidocaine. Viral vectors were delivered stereotaxically through a glass injection pipette, which was left in place for at least 5 min after injection to minimize reflux. The implantation of imaging cannulas, coverslips, or prisms was adapted to the individual experimental preparation and is described separately below. In each case, the implant and head bar were secured to the skull using dental cement for subsequent in vivo imaging.

#### Cortical windows

Prism-based imaging of cortical layer 5 pyramidal neurons was performed in Rbp4-Cre mice. To induce expression of GCaMP8s in layer 5 pyramidal neurons, an injection of adeno-associated virus (AAV) carrying the Cre-dependent calcium indicator construct ssAAV-9/2-hSyn1-dlox-jGCaMP8s(rev)-dlox-WPRE-SV40p(A) was made in the somatosensory cortex (Viral Vector Facility, University of Zurich, identifier v627-9, titer 6.8×1012 vg/ml, diluted 1:1 with Ringer’s solution). Viral injections were performed at two sites: anterior-posterior (AP) -0.7 mm; medial-lateral (ML) +3 mm from bregma and dorsal-ventral (DV) -0.6 mm from the surface of the dura; and AP -1.1 mm; ML +2.4 mm, and DV -0.6 mm. A craniotomy was then made over the primary somatosensory cortex with a 3 mm biopsy punch. To guide prism insertion, a razor blade was slowly lowered into the brain to a depth of ∼1mm. The prism assembly was composed of a reflective glass prism (1 mm, aluminum coated, N-BK7, Edmund Optics) attached to a double window (composed of a 3- and 4-mm glass coverslip attached with UV-curable optical glue) with UV-curable optical glue (NOA81, Norland). The prism was then implanted with the flush side parallel to the medio-lateral axis and facing rostrally, such that the prism provided optical access to layer 5 somata and their dendritic arbors across cortical depth.

Glutamate imaging was performed in Rbp4-Cre mice using an intersectional Cre/Flp-dependent expression strategy to semi-sparsely express the glutamate sensor iGluSnFR4s (ref. 47). A Flp-dependent iGluSnFR4s viral construct (ssAAV-1/2-dFRT-iGluSnFR4s; Viral Vector Facility, University of Zurich, identifier v1257-1, titer 7.8×1012 vg/ml) was combined with a Cre-dependent Flp-EBFP construct (ssAAV-1/2-Dlox-FlpO-EBFP; identifier v175-1, diluted to 2.85 x1012 vg/ml with sterile saline). The diluted Dlox-FlpO-EBFP and DFRT-iGluSnFR4s preparations were subsequently mixed at a 1:1 volume ratio. The viral mixture was injected into the posterior parietal cortex at AP -2.0 mm, ML +1.6 mm from bregma, and DV -0.4 mm from the surface of the dura. In Rbp4-Cre-positive layer 5 pyramidal neurons, Cre-dependent FlpO expression enabled recombination and expression of the Flp-dependent iGluSnFR4s construct. A 3 mm craniotomy was then performed over the injection site and a double window (composed of a 3- and 4-mm glass coverslip, as above) was implanted above the target region.

#### Hippocampal windows

For hippocampal calcium imaging, wild-type C57BL/6 mice were injected with ssAAV-9/2-mCaMKIIα-jGCaMP8m-WPRE-bGHp(A) (Viral Vector Facility, University of Zurich, identifier v630-9, titer 1.0 × 1013 vg/ml) in hippocampal CA1 (coordinates: AP -2.0 mm; ML -1.5 mm from bregma, and DV - 1.3 mm from the surface of the dura). Following an expression period of 2 weeks, a 3-mm-diameter craniotomy was made above the target hippocampal region56,57. For attachment, two layers of light-curing adhesive (Scotchbond Universal, 3M) were applied to the skull, followed by a ring of dental cement (Charisma, Kulzer) to prevent overgrowth with skin. A 3-mm diameter biopsy punch (BP-30F, KAI) was inserted until it reached the corpus callosum and left in place for 5 min. With a flatly cut-off injection cannula (Sterican 27G, Braun) connected to a vacuum pump, the cortex was removed until the corpus callosum became visible. Bleeding was stopped with absorbent swabs (Sugi, Kettenbach) and hemostatic sponges (Spongostan, Ethicon). Then, a cylindrical metal cannula (diameter = 3 mm and height = 1.3 mm) attached with dental cement to a 0.17-mm thick coverslip (diameter = 3 mm) was inserted into the cavity and fine-positioned with a glass capillary attached to the stereotaxic frame. The hippocampal window was fixed in place using dental cement (Tetric EvoFlow, Ivoclar). Finally, tissue glue (Vetbond, 3M) was used to connect the animal’s skin with the ring of dental cement. A head bar was attached to the Charisma ring using dental cement (Tetric EvoFlow). For imaging in CA2/CA3 (Figure 4), the window was positioned approximately 0.2 mm more laterally, and also the imaging location was at the lateral edge of the window itself. For imaging across CA1-CA3 using the scanning-space configuration (Figure S6), expression of the red-shifted calcium indicator R-CaMP1.07 (ref. 58) was virally induced (ssAAV-9/2-hSyn1-RCaMP1.07; Viral Vector Facility, University of Zurich, identifier v224-9, titer 1.5e12vg/mL) at two injections sites in hippocampal CA1 (injection coordinates: AP - 2.0 mm, ML -1.5 mm from bregma, and DV -1.3 mm from the dura) and CA3 (AP -1.7 mm, ML -2.0 mm, and DV -1.3 mm).

For prism-based hippocampal imaging (Figure S8), Thy1-GCaMP6f transgenic mice were used. Procedures followed the protocol described above for the conventional hippocampal window. A reflective glass prism (0.5 mm, aluminum coated, N-BK7, Edmund Optics) attached to a 3-mm coverslip and cannula assembly was positioned adjacent to the CA1 pyramidal-cell layer, with the side of prism viewing laterally, following existing procedures39,40. The prism thus provided an approximately sagittal optical view of CA1 somata and their apical dendrites.

#### Cerebellar windows

Purkinje cell calcium imaging was performed in wild-type mice injected with ssAAV-PHP.eB/2-L7-6-jGCaMP8m-WPRE-bGHp(A) (Viral Vector Facility, University of Zurich, identifier v968-PHP.eB). The L7-6 promoter was used to restrict indicator expression to cerebellar Purkinje cells. Before injection, the viral preparation was diluted 1:1 with Ringer’s solution. A 3-mm diameter craniotomy was centred approximately at the planned injection sites. Approximately 600 nl of virus was injected at a depth of 300 µm from the dura, at -1.2 mm AP from lambda and split between +1.2 mm and +1.5 mm ML from the midline. An imaging cannula (diameter 3 mm, height 1.3 mm) fitted with a 3-mm glass coverslip was positioned over the exposed cerebellar surface and secured to the skull using dental cement.

### Microscopy setups

For the infinity-space configuration, we used a custom-built resonant-scanning two-photon microscope as described before56. A femtosecond-pulsed laser (MaiTai, Spectra-Physics; 905 nm) was sent through an electrically tunable lens (ETL; Optotune), a scan engine consisting of an 8-kHz resonant scanner (Cambridge Technology), a 2x magnifying relay lens system and a slow galvo scanner (Cambridge Technology, 6215H), scan lens (Sill Optics, S4LFT0089/98) and tube lens (Ploessl lens consisting of two 400-mm focal length achromatic doublet lenses; AC508-400-AB, Thorlabs) before entering the objective’s back aperture. A 10x long-working-distance water-immersion objective was used across all experiments (XLPLN10XSVMP Olympus, NA 0.6, WD 8 mm), resulting in a large FOV (600-1000 µm). Fluorescence emitted from the sample was separated from the excitation path by a dichroic mirror and directed through the collection optics onto a photomultiplier tube (H10770-40, Hamamatsu). A K-mirror attached to a rotation mount was inserted into the collimated beam path between the tube lens and the objective. Positioning the K-mirror in this section of the microscope allowed rotation of the scan field relative to the specimen without rotating the sample or the scanner assembly. Mechanical rotation of the K-mirror by an angle θ produced a 2θ rotation of the field of view. K-mirrors introduced an additional folded optical path described in Figure 2. Because the beam was collimated between the tube lens and objective, this additional path did not substantially change beam convergence. The ETL was positioned in the excitation beam path upstream of the scan engine in a position optically conjugate with the objective’s back focal plane. The ETL was used to control the axial focal position without mechanical movement of the objective or specimen. The ETL was driven with waveforms synchronized to the slow galvanometric scan axis, resulting in tilted or curved imaging surfaces within single frames. For multiplane acquisition, the within-plane curvature was superimposed onto the axial separation between planes. The microscope and the ETL were controlled with ScanImage54, with a custom modification for within-plane z-scanning described below.

We implemented the scanning-space configuration in another custom-built resonant scanning two-photon microscope. The laser beam (MaiTai DeepSee, Spectra-Physics; 980 nm) was magnified by a telescope consisting of 25 mm and 60 mm focal length lenses, then sent through an electrically tunable lens (ETL; Optotune), a telescope consisting of 150 mm and 75 mm focal length lenses, and a resonant-galvo scan mirror pair (8-kHz resonant scanner; Thorlabs), before entering scan lens (100 mm), tube lens (375 mm), and a 16x objective (N16XLWD-PFNA 0.80, WD 3.0 mm; Nikon). A large-aperture commercial K-mirror (IRMU-FD30; ScienceEdge) was inserted between scan lens and tube lens, with the distance between scan lens and the entrance of the K-mirror box set to 129 mm. This distance was necessary to enable coupling-in of an additional laser beam for optogenetic stimulation (not used for this study). The physical distance between scan lens and tube lens was reduced by the optical path introduced by the K-mirror.

### Custom code for within-plane z-scanning

Custom code to control within-plane z-scanning on arbitrary scan paths and detailed instructions how this code can be inserted in a standard version of the two-photon microscopy control software ScanImage54 is openly provided via GitHub (https://github.com/HelmchenLabSoftware/K-mirror). Briefly, the code consists of a MATLAB graphical user interface that is independent of ScanImage and which allows the user to interactively choose an arbitrary z-scan trajectory for within-plane scanning. This trajectory is saved to disk. The second component is code to be inserted in the openly accessible code part of ScanImage, which loads the trajectory from disk and uses it for within-plane z-scanning. An existing checkbox of the ScanImage interface is used to flexibly enable and disable within-plane scanning.

### Design and fabrication of the 3D-printed K-mirror holder

The custom K-mirror holder was designed as a monolithic 3D printed backbone to ensure structural rigidity and optical alignment (Supplementary Movie 4). The device holds three square mirrors of 50.8 x 50.8 x 6.0 mm. To enable fine alignment of the optical path, the central mirror was mounted on an integrated flexure mechanism. Fine angular tuning of this mirror was achieved via two independent adjustment screws (XBMF3 precision screw, MISUMI). During initial prototyping, the mirrors were temporarily secured within the holder using spring-loaded M6 plunger screws threaded into dedicated bores (Delrin (POM) BN13378 1551426, Bossard). For final use, these temporary fasteners were removed to avoid stress-induced mirror curvatures, and the mirrors were permanently bonded into position using an RTV silicone adhesive (Liqui Moly #6184). The holder was printed using an SLS printer (Formiga P110, EOS) utilizing Polyamide 12 (PA2200, EOS). The printing parameters were set to a layer thickness of 100 µm with a laser spot diameter of 400 µm. During post-processing, the surface was smoothed chemically using an SF50 machine (AMT). The K-mirror holder was integrated into the microscopy beam path by mounting it onto a 2-inch rotation mount (LCRM2A/M, Thorlabs). The mechanical connection was achieved using four 3D-printed holes fitted with press-fit brass threaded inserts (Bossard) to attach the K-mirror to the rotation mount using M5 screws. The 3D-files in STEP and STL format along with a list of materials are openly available in the project’s GitHub repository (https://github.com/HelmchenLabSoftware/K-mirror).

The Leica K-mirror was mounted in a 60-mm cage system (2x plates LCP01/M, 2x rods L = 150 mm 6-mm diameter; Thorlabs) with custom 3D printed fitting rings (2x, ID 47 mm, OD 51 mm, L = 10 mm, material: PLA).

### Two-photon imaging

Laser power, zoom level and lines per imaging frame - which determines frame rate - were adjusted according to the brain structure of interest and to the expected speed of fluorescent events. Laser power ranged from 20 to 100 mW, depending on the imaging depth and preparation. In our experiments, depth-adjustment of laser power within planes was not applied, but we provide a ScanImage compatible GUI via GitHub that enables such adjustments (included in https://github.com/HelmchenLabSoftware/K-mirror). For hippocampal and cortical population imaging, single-plane recordings were acquired at 30 Hz using an image format of 512 x 512 pixels, or at 15 Hz at 1024 x 1024 pixels. For tilted or curved imaging using an ETL, the focal position was varied continuously as a function of the slow-scan axis. The ETL command voltage was interactively adjusted using a custom GUI (described below).

For cerebellar Purkinje cell imaging, z-stacks were first acquired with the slow stage motors to determine the three-dimensional arrangement of the Purkinje-cell somatic layer and the dendritic arbors. The z-stack was acquired across a range of approximately 120 µm. Then, functional recordings were performed with curved imaging planes aligned with the structures of interest in a single imaging plane (Figure 5F) or with the same curved imaging plane repeated in two spatially spaced imaging planes to simultaneously follow the superficial dendrites and the deeper somatic layer (Figure 5D). Recordings were acquired at approximately 15 Hz per plane, using an image format of 1024 x 1024 pixels.

For glutamate imaging, overview images (512 x 512 pixels) were acquired to identify dendritic processes and to determine the orientation of the dendrite selected for high-speed imaging. The field of view was subsequently rotated using the K-mirror to align the long axis of the selected dendrite with the restricted scan field. High-speed glutamate recordings were acquired using a raster scan containing 48 lines along the slow scan axis and 512 pixels along the resonant fast axis, resulting in an effective frame rate of 316.8 Hz. For high-speed glutamate imaging, no within-frame axial modulation was applied.

### Optical characterization

To assess beam pointing stability of K-mirror devices, we aligned a collimated continuous-wave laser emitting at 635 nm (CPS635R, Thorlabs) into a 2” cage system and projected the laser beam onto millimeter paper placed approximately *L* = 2 m from the cage system. The K-mirror with its rotational mount was then inserted into the 2” cage system, and the trajectory of the laser beam during a full 360° rotation was manually traced on the paper. Lateral displacements *x* were converted into angular deviations *γ* using simple trigonometry: *γ* = atan(*x/L*). Pointing stability of the 3D-printed K-mirror was fine-tuned using its adjustment screws (Figure 2B)

To measure light transmission of K-mirrors (Figure 2G, left), a 5 mm-diameter pulsed laser beam (MaiTai, Spectra Physics, 905 nm) was directed through the K-mirror, and laser power was measured before and after the K-mirror using a power meter. To measure excitation efficiency (Figure 2G, right), a homogeneously fluorescent plastic slide was placed below the objective, and fluorescence intensity was measured for different K-mirror configurations without repositioning the sample. Measurements were performed at low excitation power (approx. 2 mW) and short exposure times to minimize photobleaching. Excitation efficiencies were normalized to fluorescence obtained without a K-mirror. Theoretical excitation efficiencies were computed as the square of the measured transmission values.

To determine microscope resolution (Figure 2H; Figure S3), 1 µm-diameter fluorescent beads were suspended in distilled water between a glass slide and a 0.17 mm thick glass cover slip.

### Geometrical optics simulations

To quantify beam clipping and pointing errors introduced by the K-mirror, we used an AI-assisted workflow (Claude Code (Anthropic)) to build sequential ray-tracing models in Python using the open-source raytracing package Optiland (v0.6.2; ref. 59). Optical surfaces (scan and tube lenses, flat mirrors) were simulated as paraxial surfaces with the clear aperture of the respective hardware component. The K-mirror was modelled as three flat mirrors in the “K” arrangement; rotation of the assembly by an angle *φ* was simulated as a rigid-body rotation in 3D. For the scanning-space configuration, the beam was traced from the scan mirrors through the scan lens, the K-mirror housing (α = 36.6°, matching the ScienceEdge IRMU-FD30 K-mirror), and the tube lens. For the infinity-space configuration, the collimated beam was traced from the tube lens through the K-mirror (2″ flat mirrors, α = 29°) to the objective back aperture.

In the scanning-space configuration model, the input beam was defined by its 1/e2 diameter and represented as a bundle of rays sampled on hexapolar rings out to 1.5 times the 1/e2 beam radius (98.9% of the power of the Gaussian beam), with each ray weighted by exp(™2*r*2/*w*2) with radial position *r* and beam waist *w*. A ray was considered clipped when it fell outside a physical aperture. Transmission is reported as the Gaussian-power-weighted fraction of rays reaching the housing exit. In the infinity-space configuration model, the input beam was defined as a top-hat profile, reflecting that the modelled microscope used a large beam that was clipped by the scan mirrors. Both simulations are provided as interactive Jupyter notebooks on GitHub, allowing all optical and mechanical parameters to be adjusted to evaluate alternative microscope designs (https://github.com/HelmchenLabSoftware/K-mirror).

### Histology

After completion of calcium imaging experiments, animals were administered a lethal dose of pentobarbital (Ekonarcon, Streuli) and transcardially perfused with 0.1 M phosphate buffered saline (PBS) followed by 4% paraformaldehyde (in 0.1 M PBS). 150 µm thick coronal brain sections were stained with the nuclear stain DAPI and imaged with a confocal laser-scanning microscope (Leica SP8) in tile-scan mode using a 10x objective and 488 nm excitation to record GCaMP fluorescence. No antibodies were used to enhance intrinsic signals. Histology was performed in a subset of experiments to confirm localization of implants or injections.

### Preprocessing of two-photon imaging data

TIFF files recorded with ScanImage were processed using a custom MATLAB pipeline. A common PMT offset was estimated for each recording session from the 10th percentile of pixel intensities sampled across the raw image files and subtracted from all frames before image registration. Alternating scan lines were corrected for a horizontal phase offset using fast Fourier transform-based cross-correlation between subsequent lines. Phase shifts were averaged across line pairs, and the median displacement across frames was applied as a common integer-pixel shift to align odd and even scan lines.

Motion correction was performed using NoRMCorre60. First, rigid registration was applied by estimating a single global horizontal and vertical displacement for each frame relative to reference with subpixel precision. Where required, the rigidly corrected recordings were subsequently subjected to non-rigid correction to compensate for residual local deformation. Corrected recordings and their corresponding mean images were visually inspected for potential motion artifacts and to assess motion-correction quality. For glutamate imaging data, only rigid motion correction was performed.

Following motion correction, somatic and dendritic regions of interest (ROIs) were manually defined using an interactive MATLAB-based ROI-drawing toolbox (https://github.com/PTRRupprecht/Drawing-ROIs-without-GUI; ref. 61) based on anatomical overviews or activity-based overlays. For each ROI, the raw fluorescence trace F(t) was calculated by averaging the intensity of all pixels contained within the ROI. Relative fluorescence changes were subsequently calculated as ΔF/F(t) = (F(t)-F0 (t))/F0(t), with F0(t) estimated using a moving percentile filter (20 s filter size; 20th percentile for all recordings except for glutamate imaging, where a 20-s median filter was used).

### Spatial mapping of activity components and decay times

Motion-corrected calcium imaging data recorded from hippocampal CA2/3 were used for analysis of subcellular activity and calcium decay kinetics (Figure 4). Somatic ROIs were manually defined, and the corresponding fluorescence and ΔF/F traces were extracted as described above. From somatic ΔF/F traces, clearly visible events that reflect spike bursts were identified using a visually defined threshold (300% ΔF/F). For each detected event, a time window (100 frames before and 200 frames after the event, frame rate 15 Hz) of the movie was extracted and averaged to obtain event-triggered calcium transients. For pixelwise spatial mapping of these average events and their half-decay time, we computed the map of local correlations37 (correlation map in Figure 4) and used a visually determined threshold applied to this correlation map to create a binary mask for each neuron. Analysis of decay kinetics was only performed within this mask (Figure 4H,J). The half-decay time was computed as the time required for the fluorescence signal to decrease to 50% of the peak amplitude for the averaged calcium transient. The resulting half-decay values were assigned to their corresponding pixel coordinates to generate spatial maps of calcium-decay kinetics.

### Analysis of subcellular glutamate signals

Subcellular ROIs were manually drawn as described above, covering dendritic segments or individual spines. To separate global from local activity, a global trace was calculated for each ROI as the frame-by-frame median of all remaining ROI ΔF/F traces. This global trace was linearly fitted to the corresponding ROI trace, and the fitted component was subtracted to obtain the local residual.

### Denoising of calcium imaging data

For visualizations (Supplementary Movies 1 and 3), raw two-photon calcium imaging recordings were denoised using DeepCAD-RT62,63. After bidirectional alignment, separate models were trained for hippocampal (Supplementary Movie 1) and cerebellar recordings (Supplementary Movie 3) to account for differences in recording conditions. For each recording, the model was trained on the same recording that was subsequently used for inference, using the default DeepCAD-RT parameters (3D patch size: 150*150*150 pixels [x*y*t], 10 training epochs, learning rate: 5*10^−5^, training patch overlap: 0.25, and inference patch overlap: 0.6). The model from the final training epoch was used for inference. Denoised movies were used for supplementary movies, not for analysis.

### Statistics

No statistical tests were performed for this manuscript. Box plots used standard settings in MATLAB 2022b, with the central line at the median of the distribution, the box at the 25th and 75th percentiles, and the whiskers at extreme values excluding outliers.

## Supplementary Figures

**Supplementary Figure S1.**
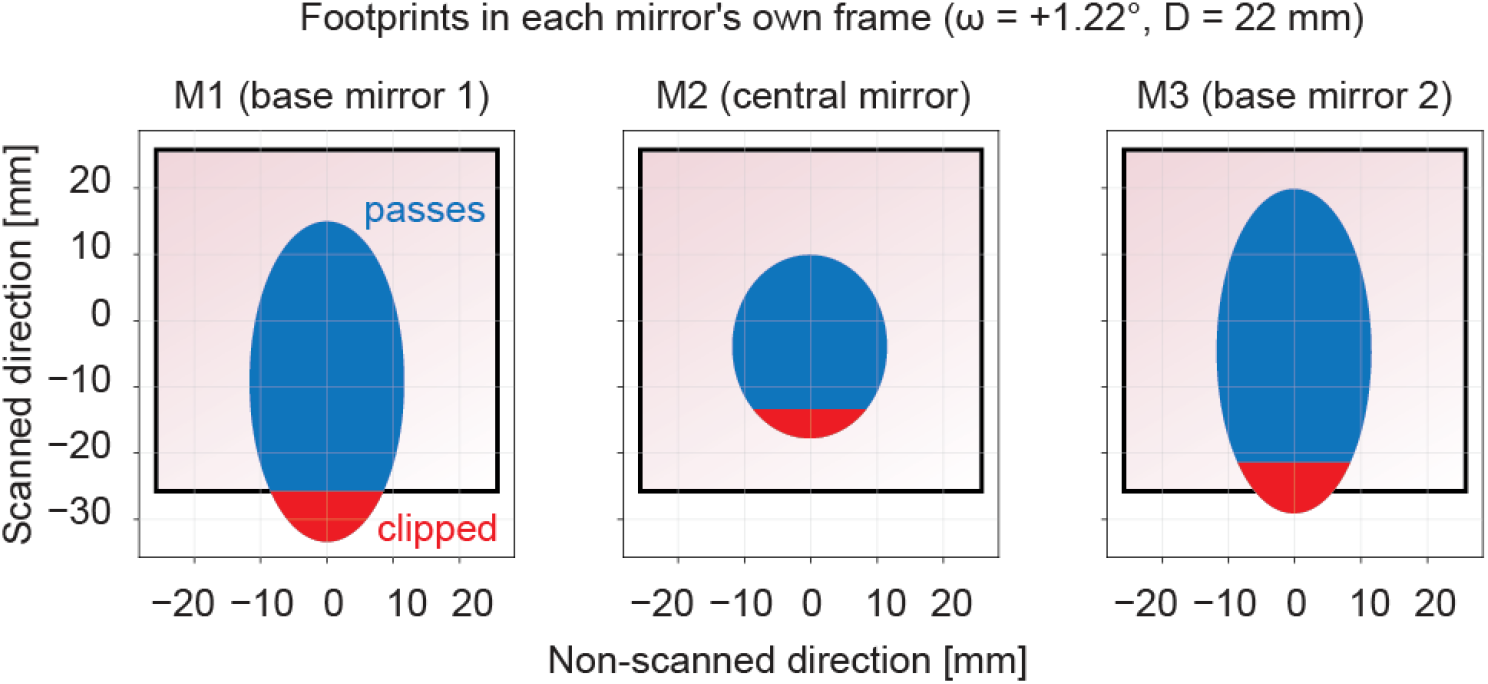
Beam clipping for angular scanning. Top-hat beam profile (beam diameter D = 22 mm) passing through the K-mirror system as described in Figure 2D (d_∞_ = 100 mm, ω = +1.22°). Normal reflections are colored in blue, beam clipping in red. Mirrors (50.8 mm sidelength) are indicated as black outlines. The footprint of the beam is elliptical on the mirrors due to their tilt. The beam is primarily clipped by base mirror 1 (M1).

**Supplementary Figure S2.**
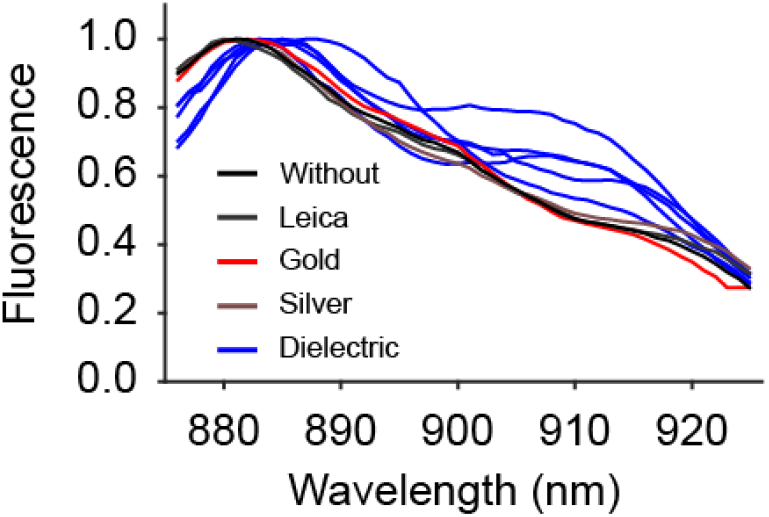
Wavelength dependent fluorescence yield for dielectric mirrors. Fluorescence yield, measured using a homogeneous fluorescence slide while changing the wavelength of the tunable laser. Without K-mirror, or with the Leica K-mirror, or with gold, or silver 3D-printed K-mirrors, the recorded spectrum simply reflects the product of laser output and the transmittance of the microscope components. For the dielectric K-mirror, the dielectric layers at shallow reflective angles act as chirped mirrors that induce temporal dispersion of laser pulse, thereby affecting fluorescence excitation efficiency in a wavelength-dependent manner. This behavior is different for the four instances of the dielectric K-mirror (rotated in increments of 45°) and therefore difficult to predict during routine operation. The clear wavelength-dependence of fluorescence yield has previously been shown to be a hallmark of undesired pulse-chirping effects of dielectric mirrors^26^, in this case likely introduced by the shallow reflecting angles within the K-mirror.

**Supplementary Figure S3.**
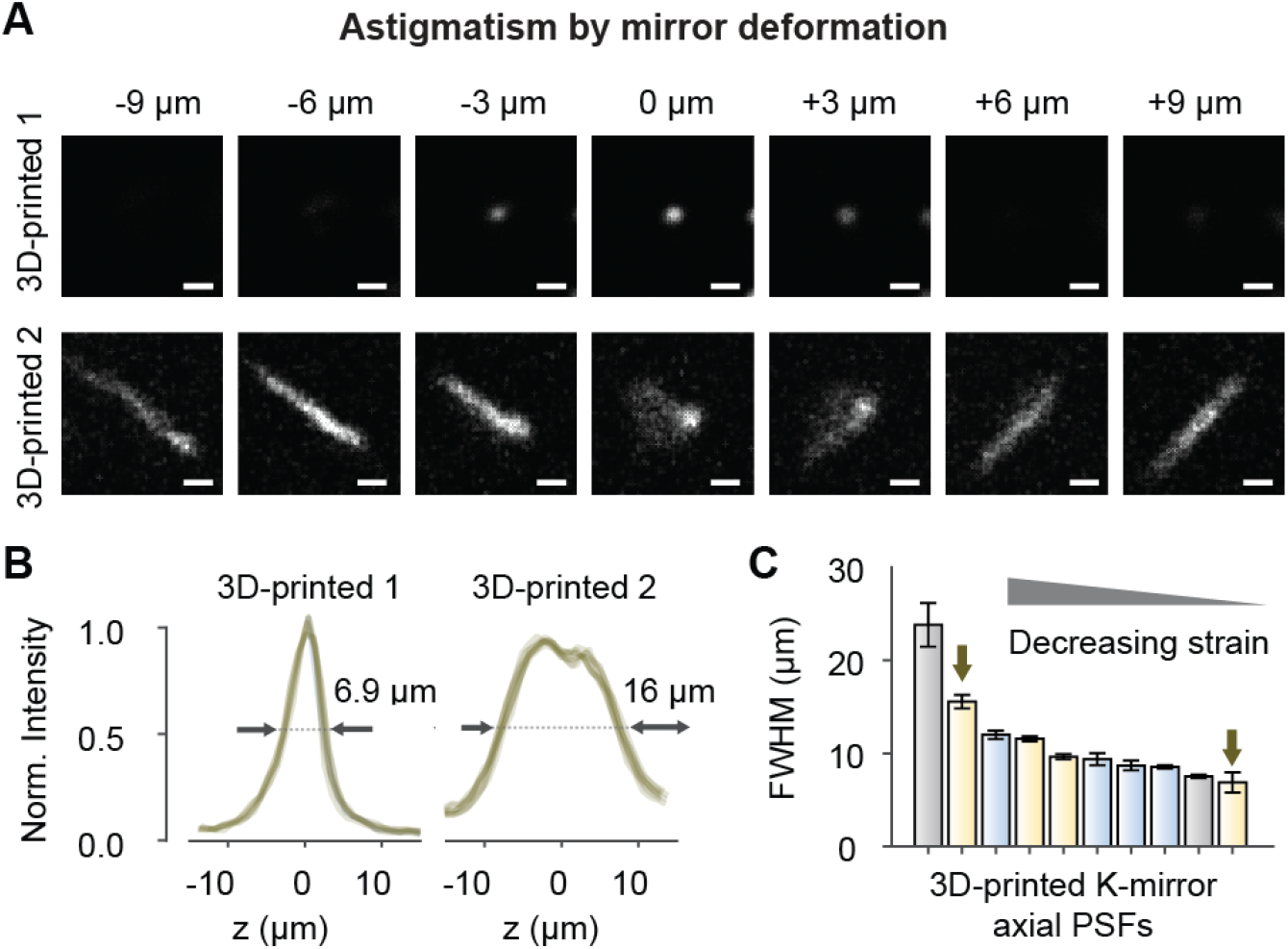
Astigmatism introduced by distorted surfaces in a K-mirror. **A**, The large mirror surfaces of the K-mirrors are highly susceptible to introducing optical aberrations if any force (e.g., clamping screws) is applied to them. For 3D-printed K-mirrors, the normal point spread function (upper row; “3D-printed 1”) degraded when spring-loaded plastic screws applied pressure to hold mirrors in place, inducing a clear astigmatim (lower row; “3D-printed 2”). Scale bars are 2 μm. **B**, Astigmatism decreases the resolution, measured as axial FWHM (examples shown for 16 μm vs. 6.9 μm with the same gold mirrors used in the 3D-printed K-mirror). **C**, With decreasing strain, FWHM improved and approached the resolution obtained without K-mirror. For the final design, low-strain silicone glue was used to replace spring-loaded screws entirely. Colors indicate mirror types (gray: silver mirrors; yellow: gold mirrors; blue: dielectric mirrors). PSFs shown in (B) are highlighted with arrows. All PSFs were measured with 1 μm-beads.

**Supplementary Figure S4.**
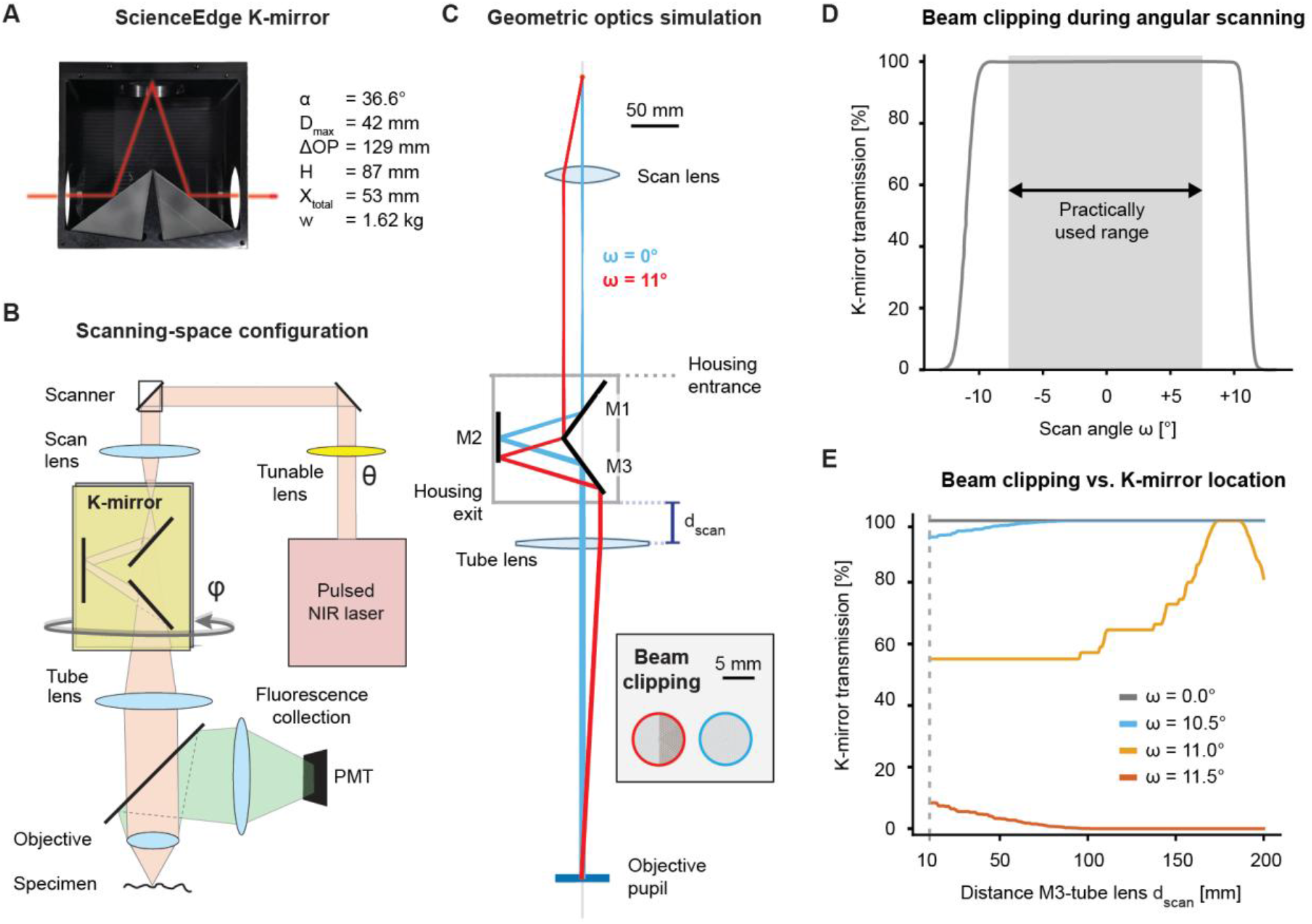
Scanning-space configuration to insert a K-mirror into a point-scanning two-photon microscope. In this configuration, the K-mirror is inserted between scan lens and tube lens, leaving room for other optical elements between tube lens and objective. **A**, A large-aperture commercial K-mirror (IRMU-FD30; ScienceEdge) was used for this configuration. With a mirror length of 70.8 mm, the maximal beam diameter is 70.8 mm · sin(36.6°) = 42.2 mm. The ScienceEdge datasheet more conservatively specifies the maximal beam diameteras37.2 mm. **B**, To compensate for the added optical path length introduced by the K-mirror (ΔOP), the physical distance between scan lens and tube lens was reduced accordingly. Since the distance between scan lens and tube lens was adapted to compensate for the optical path introduced by the K-mirror, no direct comparison with and without K-mirror is possible in this configuration with reasonable effort. **C**, Geometric optics simulations of the scanned beam. In this specific simulation, which corresponds to the real microscope used in Figure S6, the beam diameter at the scanner (top) is 1.44 mm and is expanded by the telescope consisting of scan lens (focal length 100 mm) and tube lens (375 mm). K-mirror flat mirrors are indicated as M1-M3. Beam clipping at objective back aperture is illustrated for scan angles ω = 0° (no clipping, blue) and ω = 11° (approx. 50% clipping, red), with transmitted beam shown in light grey and blocked beam shown in dark gray. The colored rings indicate the size of the unblocked beam at the objective back aperture. **D**, Beam clipping as a function of the scan angle ω. The maximum scan angle is limited by the range of the 8 kHz scanner (±13° optical). In practice, the useful scan range is lower due to the limited field of view of the objective. In our specific microscope configuration, the beam is not clipped by the K-mirror in the practically used scan range. **E**, Beam clipping changes with the position of the K-mirror assembly with respect to the tube lens (distance from K-mirror external housing to center of tube lens, indicated as *d*_*scan*_ in (C)). Beam clipping is almost negligible for scan angles up to 10.5°; for scan angles >11.5° (up to the maximum scan angle of 13°) almost the entire beam is clipped. Python notebooks to reproduce and adapt the simulation to other parameters and microscope designs are on GitHub (Methods).

**Supplementary Figure S5.**
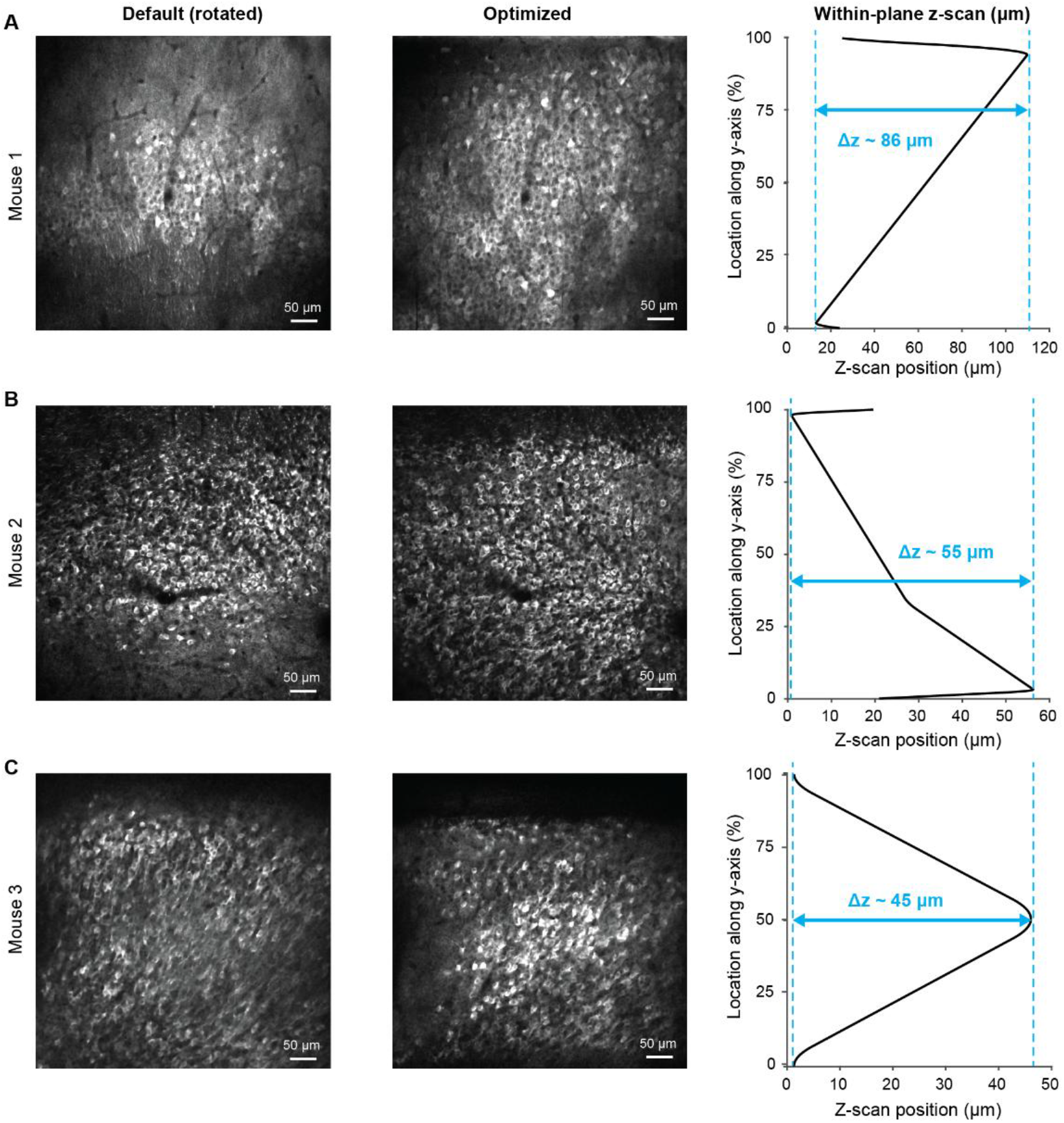
Axial scan trajectories to align the scan field with the CA1 pyramidal cell layer. *Left:* Mean fluorescence after rotation but before the application of axial scanning across 3 animals (compare Figure 3F). *Middle:* Mean fluorescence optimized by rotation and axial scanning. *Right:* Axial scan trajectories specific to each of the optimizations. For “Mouse 1” (**A**) and “Mouse 2” (**B**), a mostly linear z-scan trajectory (“tilt”) is sufficient to align the pyramidal cell layer with the scan field. For “Mouse 3” (**C**), the pyramidal cell layer matched with a curved, convex z-scan trajectory.

**Supplementary Figure S6.**
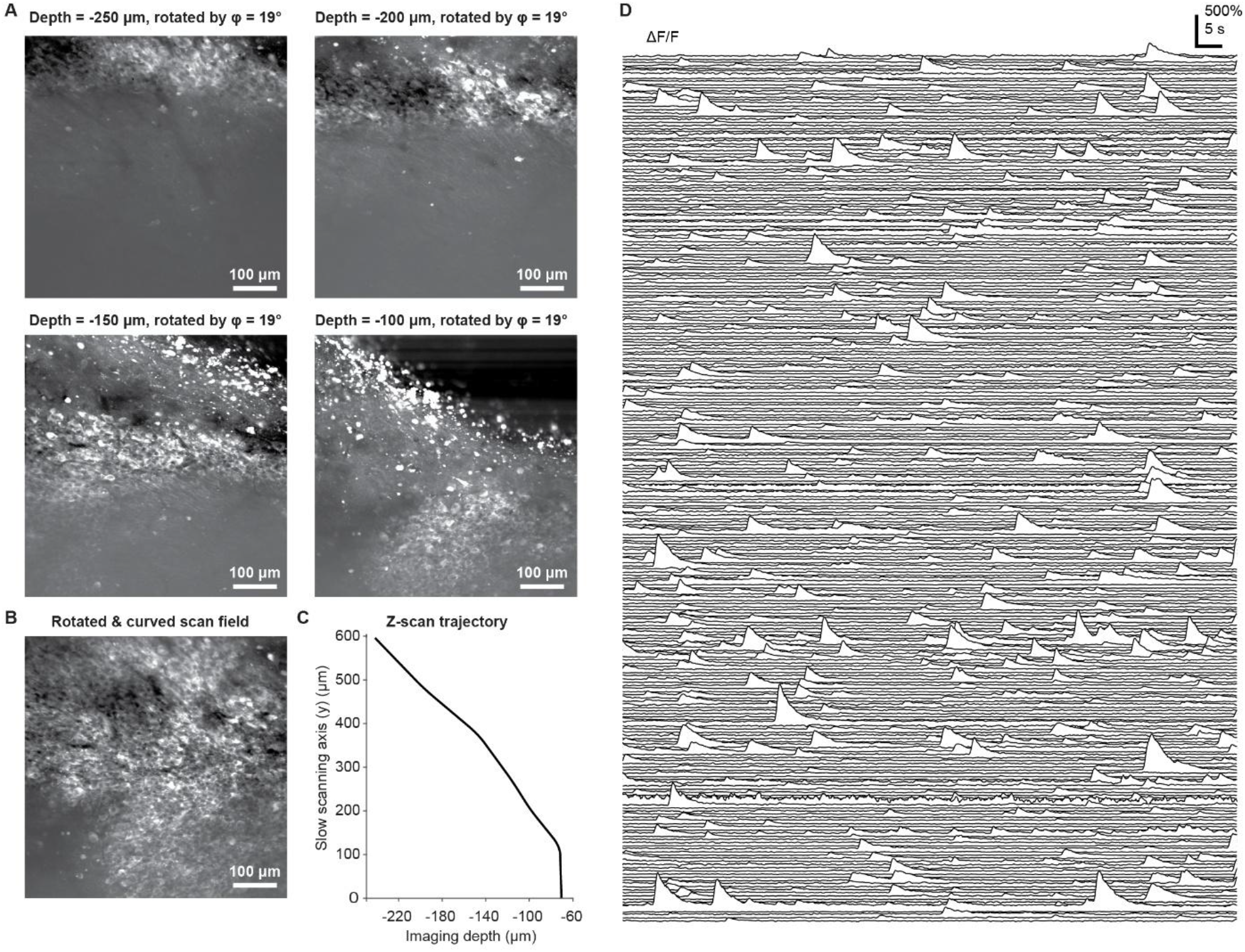
Scan field optimization for pyramidal cell layer imaging across CA1-CA3 using the scanning-space K-mirror configuration. For imaging, expression of the red-shifted calcium indicator R-CaMP1.07 was virally induced, and imaging was performed through a cannula with coverslip as before (cf. Figure 3). For the description of the scanning-space configuration, see Figure S4. **A**, Z-stack across hippocampal CA1-CA3. Using scan field rotation with the K-mirror, the FOV was rotated (*φ* = 19°) such that the curvature of the pyramidal cell layer across CA3 (top left) to CA1 (bottom right subpanel) was aligned with the slow y-axis. **B**, Scan field curvature enables imaging of pyramidal cells that can otherwise be imaged only across a z-stack (compare to panel (A)). **C**, Z-scan trajectory used to obtain the FOV in panel (B). The y-axis is aligned with panel (B). **D**, 276 simultaneously imaged CA1-CA3 neurons from the FOV in (B) (unsorted).

**Supplementary Figure S7.**
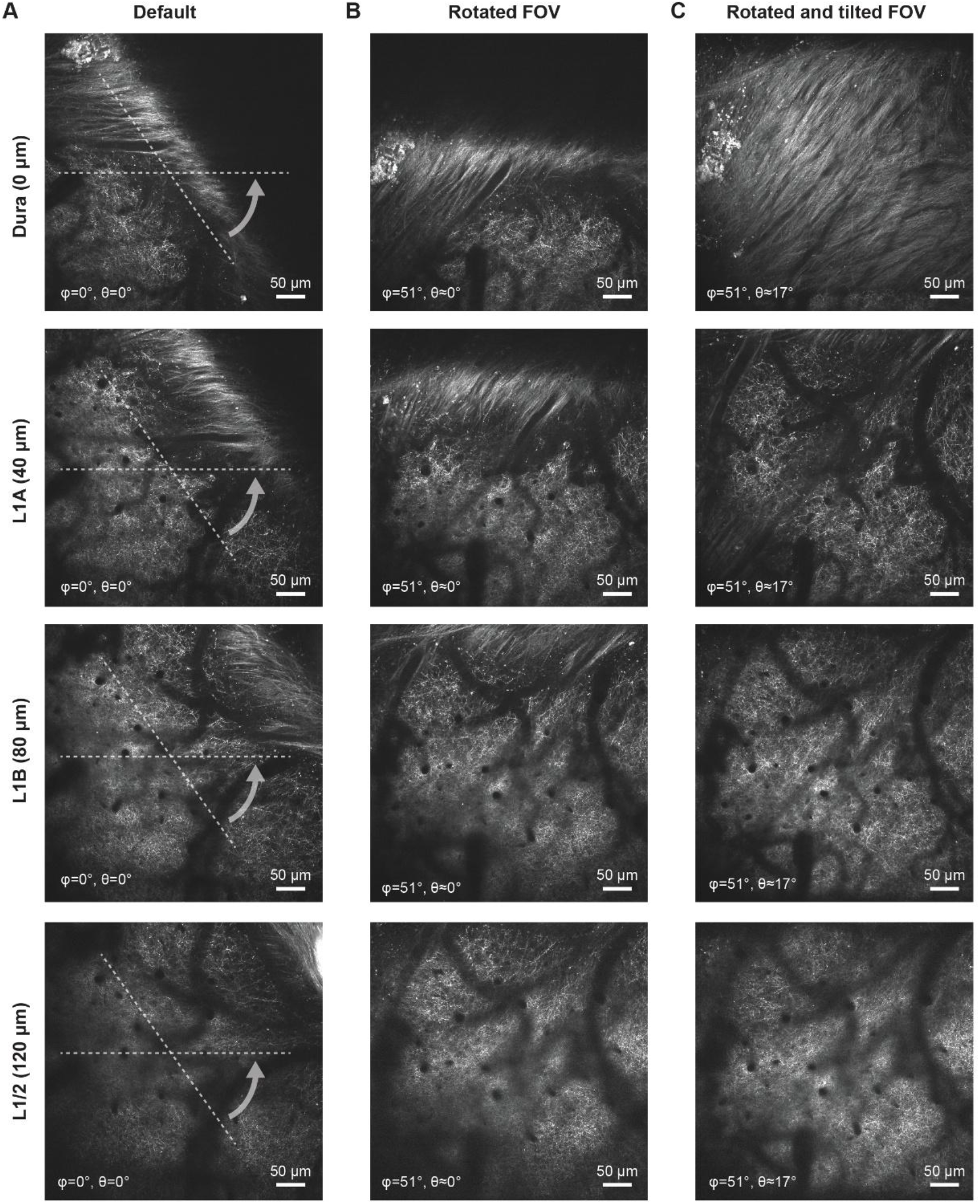
Alignment of the FOV with cortical tilt in L1 using tilted plane scanning (θ) with scan field rotation (*φ*). Axons projecting from auditory cortex to the posterior parietal cortex were labeled using tdTomato, and imaged through a 5-mm cortical window. **A**, Entire column: Default imaging without image plane optimization. **B**, Scan field rotation (*φ*) to align the tilted axis of cortical layering with the slow (y-) axis of the scanning microscope. **C**. Scan field rotation (*φ*) together with z-scanning for tilted plane imaging (θ). The dura (top image; autofluorescence) as well as superficial L1A (sparse labeling of axons) and deeper L1B (denser labeling) are visibly aligned with the scanned imaging plane.

**Supplementary Figure S8.**
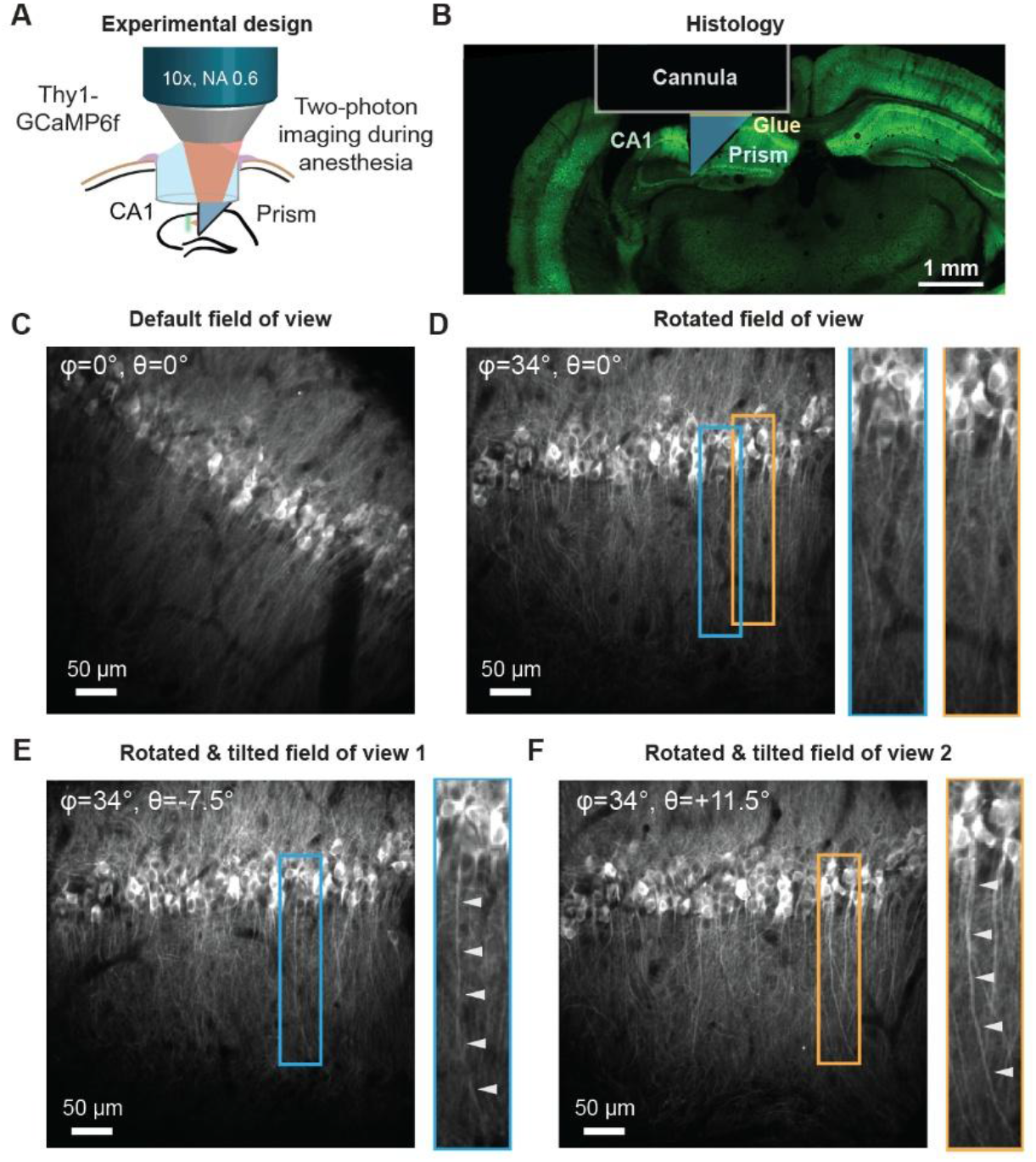
Optimized prism-based imaging of pyramidal cells in hippocampal CA1. **A**, Schematic of experimental setup. A cannula with a 3 mm-coverslip is attached with a glass prism and implanted above mouse hippocampal CA1. **B**, Coronal confocal histology image to illustrate the location and orientation of the prism with respect to hippocampal anatomy. Green is fluorescence from transgenically expressed GCaMP6f. **C**, Imaging through the prism in anesthetized animals provides an approximately para-sagittal section of CA1. **D**, After rotation of the FOV. Selected neurons are highlighted (blue and orange), with their dendrites branching out of the imaging plane. **E&F**, The rotated image plane is tilted using the tunable lens to capture the apical dendrite of specific neurons (arrowheads in zoom-ins). In (E) and (F), the axial focus is linearly shifted along the slow scanning axis with a tilt angle of θ = -7.5° (maximum axial shift of -40 μm) and of θ = +11.5° (maximum axial shift of +60 μm).

**Supplementary Figure S9.**
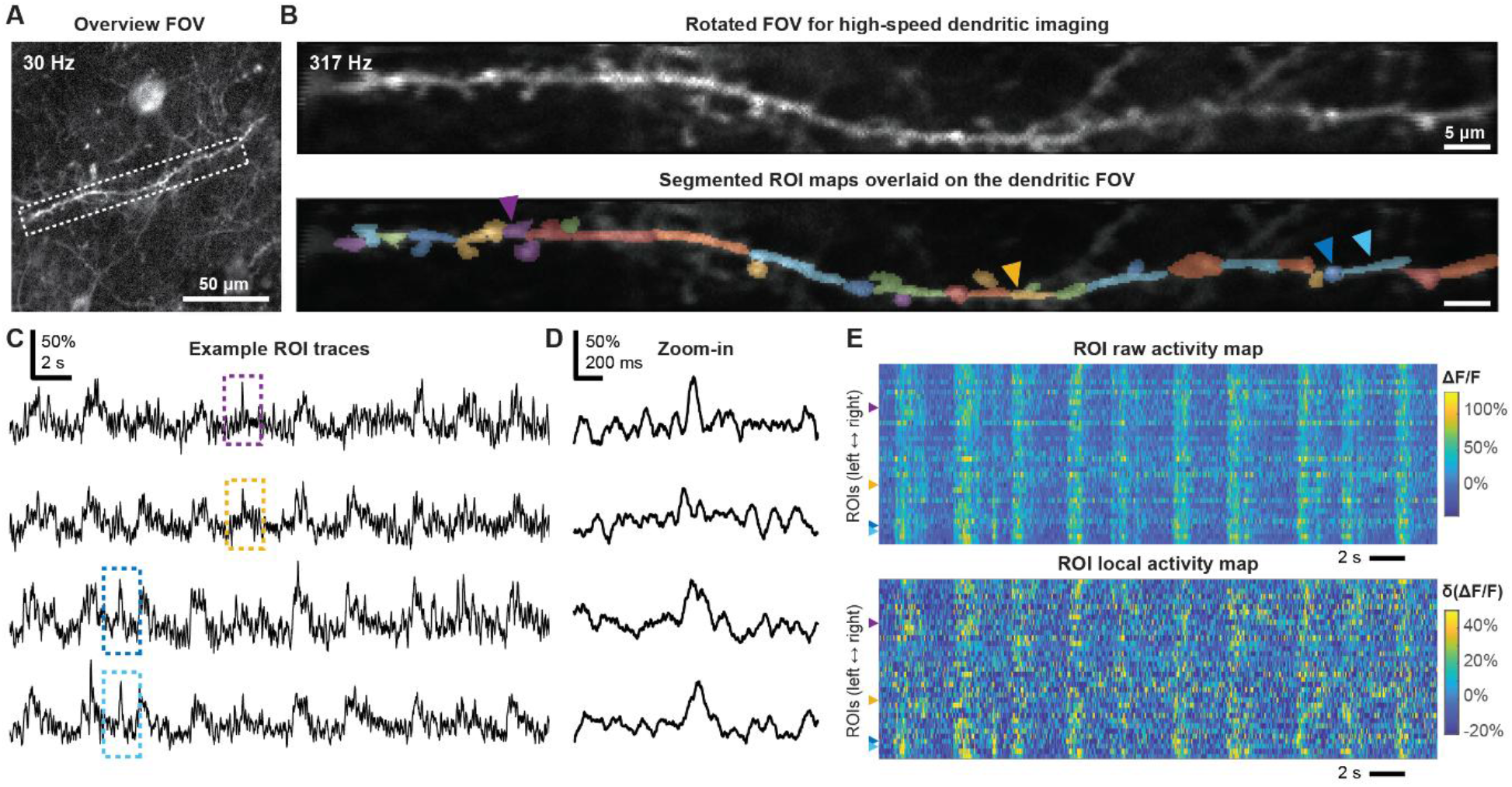
Scan field rotation for high-speed dendritic glutamate imaging, additional data from a second mouse. **A**, Imaging of iGluSnFR4s-expressing layer 5 pyramidal neurons in a mouse during anesthesia, overview with slow scanning (30 Hz, 512×512 pixels). **B**, *Top:* Rotated FOV of the dendritic segment highlighted with a dashed rectangle in (A), recorded at high speed (317 Hz, 512×48 pixels), averaged over 30,000 frames. *Bottom:* Colors indicate different ROIs from the dendritic process, covering both dendritic segments and spines. **C**, ΔF/F traces from four example ROIs (marked with arrows in (B)). **D**, Zoom-in to selected fast events. These events do not occur globally across all ROIs. **E**, Raster plots of all the ROIs ordered from left to right along the dendritic process in (B). *Top:* Raw ΔF/F of all ROIs. *Bottom:* ΔF/F after subtracting the regressed global fluorescence component, highlighting local activity.

## Supplementary Movie Legends

**Supplementary Movie 1. Scan field optimization for imaging of hippocampal pyramidal neurons**. Movie related to Figure 3. Pyramidal neurons expressing the calcium indicator GCaMP8m are imaged in the CA1 pyramidal layer of an anesthetized mouse. Three steps towards the optimization of the scan field are shown: the “default imaging plane” shows the field of view without scan field rotation or tilt; the “rotated imaging plane” shows the field of view after scan field rotation such that the curvature of the pyramidal cell layer aligns with the slow y-axis, but without adapting the curvature of the scan field to the cell layer; the “optimized imaging plane” shows the field of view after scan field rotation and alignment of the scan field curvature along the y-axis with the curvature of the pyramidal cell layer. For the video, the original movie (1024×1024 pixels) was spatially downsampled (512×512). The original calcium imaging data were recorded at 15 Hz. Raw data were denoised using DeepCAD-RT (for visualization only; Methods). Compared to real-time, the movie is sped up by 13x.

**Supplementary Movie 2. Scan field optimization for dendritic imaging of hippocampal pyramidal neurons**. Movie related to Figure 4. Pyramidal neurons expressing the calcium indicator GCaMP8m are imaged in the CA2/3 pyramidal layer of an anesthetized mouse. Three steps towards the optimization of the scan field are shown: the “default imaging plane” shows the field of view without scan field rotation or tilt; the “rotated imaging plane” shows the field of view after scan field rotation such that the apical dendrites of pyramidal cells align with the slow y-axis, but without adapting the curvature of the scan field to the dendrites; the “rotated and curved imaging plane” shows the field of view after scan field rotation and alignment of the scan field curvature along the y-axis with the orientation of pyramidal dendrites. The original calcium imaging data were recorded at 15 Hz and were smoothed with a Gaussian kernel with a standard deviation of 6 frames in time. Compared to real-time, the movie is sped up by 12x.

**Supplementary Movie 3. Somato-dendritic imaging in cerebellar Purkinje cells**. Movie related to Figure 5. Cerebellar Purkinje cells expressing the calcium indicator GCaMP8m are imaged with the imaging plane aligned to capture both dendrites and somata of Purkinje cells. Purkinje cell dendrites exhibit large, fast and frequent calcium transients, reflecting putative climbing fiber-induced complex spikes. The original calcium imaging data were recorded at 15 Hz. Raw data were denoised using DeepCAD-RT (for visualization only; Methods). Compared to real-time, the movie is sped up by 4x.

**Supplementary Movie 4. Image rotation with the 3D-printed K-mirror**. Illustration of image rotation by rotation of the K-mirror. The K-mirror is manually rotated while filmed from above. The inset shows the view through the K-mirror, with a reference object (letters “UZH”) shown.

## References

1. Schoenfeld, G., Carta, S., Rupprecht, P., Ayaz, A. & Helmchen, F. In Vivo Calcium Imaging of CA3 Pyramidal Neuron Populations in Adult Mouse Hippocampus. eNeuro 8, (2021).

2. Dong, C., Madar, A. D. & Sheffield, M. E. J. Distinct place cell dynamics in CA1 and CA3 encode experience in new environments. Nat Commun 12, 2977 (2021).

3. Rashid, S. K. et al. The dendritic spatial code: branch-specific place tuning and its experience-dependent decoupling. bioRxiv. Preprint at 10.1101/2020.01.24.916643 (2020).

4. Kveim, V. A. et al. Divergent recruitment of developmentally defined neuronal ensembles supports memory dynamics. Science 385, eadk0997 (2024).

5. Ji, N., Freeman, J. & Smith, S. L. Technologies for imaging neural activity in large volumes. Nat Neurosci 19, 1154–1164 (2016).

6. Abdelfattah, A. et al. Neurophotonic Tools for Microscopic Measurements and Manipulation: Status Report. Neurophoton. 9, (2022).

7. Denk, W., Strickler, J. H. & Webb, W. W. Two-Photon Laser Scanning Fluorescence Microscopy. Science 248, 73–76 (1990).

8. Tsien, R. Y. & Bacskai, B. J. Video-Rate Confocal Microscopy. in Handbook of Biological Confocal Microscopy (ed. Pawley, J. B.) 459–478 (Springer US, Boston, MA, 1995). doi:10.1007/978-1-4757-5348-6_29.

9. Fan, G. Y. et al. Video-Rate Scanning Two-Photon Excitation Fluorescence Microscopy and Ratio Imaging with Cameleons. Biophysical Journal 76, 2412–2420 (1999).

10. Göbel, W., Kampa, B. M. & Helmchen, F. Imaging cellular network dynamics in three dimensions using fast 3D laser scanning. Nat Methods 4, 73–79 (2007).

11. Grewe, B. F., Voigt, F. F., Hoff, M. van ‘t & Helmchen, F. Fast two-layer two-photon imaging of neuronal cell populations using an electrically tunable lens. Biomed. Opt. Express, 2, 2035–2046 (2011).

12. Botcherby, E. J. et al. Aberration-free three-dimensional multiphoton imaging of neuronal activity at kHz rates. Proc. Natl. Acad. Sci. U.S.A. 109, 2919–2924 (2012).

13. Sofroniew, N. J., Flickinger, D., King, J. & Svoboda, K. A large field of view two-photon mesoscope with subcellular resolution for in vivo imaging. eLife 5, e14472 (2016).

14. Rupprecht, P., Prendergast, A., Wyart, C. & Friedrich, R. W. Remote z-scanning with a macroscopic voice coil motor for fast 3D multiphoton laser scanning microscopy. Biomed. Opt. Express, 7, 1656–1671 (2016).

15. Duan, X., Li, H., Li, X., Oldham, K. R. & Wang, T. D. Axial beam scanning in multiphoton microscopy with MEMS-based actuator. Opt. Express 25, 2195 (2017).

16. Peinado, A., Bendek, E., Yokoyama, S. & Poskanzer, K. E. Deformable mirror-based axial scanning for two-photon mammalian brain imaging. Neurophoton. 8, (2021).

17. Katona, G. et al. Fast two-photon in vivo imaging with three-dimensional random-access scanning in large tissue volumes. Nat Methods 9, 201–208 (2012).

18. Griffiths, V. A. et al. Real-time 3D movement correction for two-photon imaging in behaving animals. Nature Methods 1–8 (2020) doi:10.1038/s41592-020-0851-7.

19. Berlage, C. et al. High-speed three-dimensional random access scanning with a linear SLM. Optica 11, 1639 (2024).

20. Naumann, H. Spiegelnde Prismen. in Optische Instrumente /Optical Instruments (ed. Flügge, S.) vol. 5 /29 211–226 (Springer Berlin Heidelberg, Berlin, Heidelberg, 1967).

21. Bretz, R. Techniques of Television Production. (McGraw-Hill Book Company, New York, 1962).

22. Hildebrand, R. H., Dragovan, M. & Novak, G. Detection of submillimeter polarization in the Orion nebula. ApJ 284, L51 (1984).

23. Byers, P. et al. Super-resolution upgrade for deep tissue imaging featuring simple implementation. Nat Commun 16, 5386 (2025).

24. Chen, B. et al. Resolution doubling in light-sheet microscopy via oblique plane structured illumination. Nat Methods 19, 1419–1426 (2022).

25. Sullivan, D. L. Alignment of Rotational Prisms. Appl. Opt. 11, 2028 (1972).

26. Rupprecht, P. Pulse dispersion, dielectric mirrors and fluorescence yield: A case report of a two-photon microscope. https://zenodo.org/record/2592465 (2019) doi:10.5281/ZENODO.2592465.

27. Göbel, W. & Helmchen, F. New Angles on Neuronal Dendrites In Vivo. Journal of Neurophysiology 98, 3770–3779 (2007).

28. O’Hare, J. K., Wang, J., Shala, M. D., Polleux, F. & Losonczy, A. Distal tuft dendrites predict properties of new hippocampal place fields. Neuron 113, 1969-1982.e7 (2025).

29. Gonzalez, K. C. et al. Synaptic basis of feature selectivity in hippocampal neurons. Nature 637, 1152–1160 (2025).

30. Kerlin, A. et al. Functional clustering of dendritic activity during decision-making. eLife 8, e46966 (2019).

31. Schoenfeld, G. et al. Unsigned temporal difference errors in cortical L5 dendrites during learning. bioRxiv. Preprint at 10.1101/2021.12.28.474360 (2024).

32. Francioni, V. et al. Vectorized instructive signals in cortical dendrites. Nature 652, 1254–1263 (2026).

33. Moore, J. J. et al. Sub-cellular population imaging tools reveal stable apical dendrites in hippocampal area CA3. Nat Commun 16, 1119 (2025).

34. Balind, S. R. et al. Distinct types of dendritic Ca^2+^ spikes in CA3 pyramidal neurons associated with neuronal activity changes during navigation. Sci. Adv. 12, eaec6015 (2026).

35. Hainmueller, T. & Bartos, M. Parallel emergence of stable and dynamic memory engrams in the hippocampus. Nature 558, 292–296 (2018).

36. Holthoff, K., Tsay, D. & Yuste, R. Calcium Dynamics of Spines Depend on Their Dendritic Location. Neuron 33, 425–437 (2002).

37. Junek, S., Chen, T.-W., Alevra, M. & Schild, D. Activity Correlation Imaging: Visualizing Function and Structure of Neuronal Populations. Biophysical Journal 96, 3801–3809 (2009).

38. Jaffe, D. B. et al. A model for dendritic Ca2+ accumulation in hippocampal pyramidal neurons based on fluorescence imaging measurements. Journal of Neurophysiology 71, 1065–1077 (1994).

39. Redman, W. T. et al. Long-term transverse imaging of the hippocampus with glass microperiscopes. eLife 11, e75391 (2022).

40. Lee, B. H. et al. Fast dendritic excitations primarily mediate back-propagation in CA1 pyramidal neurons during behavior. bioRxiv. Preprint at 10.64898/2026.01.03.696606 (2026).

41. Wu, X. et al. A dendrite-resolved, in vivo transfer function from spike patterns to dendritic Ca^2+^. bioRxiv. Preprint at 10.64898/2026.01.18.700189 (2026).

42. Tank, D. W., Sugimori, M., Connor, J. A. & Llinás, R. R. Spatially Resolved Calcium Dynamics of Mammalian Purkinje Cells in Cerebellar Slice. Science 242, 773–777 (1988).

43. Miyakawa, H., Lev-Ram, V., Lasser-Ross, N. & Ross, W. N. Calcium transients evoked by climbing fiber and parallel fiber synaptic inputs in guinea pig cerebellar Purkinje neurons. Journal of Neurophysiology 68, 1178–1189 (1992).

44. Kitamura, K. & Hausser, M. Dendritic Calcium Signaling Triggered by Spontaneous and Sensory-Evoked Climbing Fiber Input to Cerebellar Purkinje Cells In Vivo. Journal of Neuroscience 31, 10847–10858 (2011).

45. Ramirez, J. E. & Stell, B. M. Calcium Imaging Reveals Coordinated Simple Spike Pauses in Populations of Cerebellar Purkinje Cells. Cell Reports 17, 3125–3132 (2016).

46. McDonald, A. J. et al. A versatile, positive-going voltage indicator that enables accessible two-photon recordings in vivo. bioRxiv. Preprint at 10.64898/2026.04.07.717088 (2026).

47. Aggarwal, A. et al. Glutamate indicators with increased sensitivity and tailored deactivation rates. Nat Methods 23, 417–425 (2026).

48. Shih, A. Y. et al. Two-Photon Microscopy as a Tool to Study Blood Flow and Neurovascular Coupling in the Rodent Brain. J Cereb Blood Flow Metab 32, 1277–1309 (2012).

49. Kleinfeld, D., Mitra, P. P., Helmchen, F. & Denk, W. Fluctuations and stimulus-induced changes in blood flow observed in individual capillaries in layers 2 through 4 of rat neocortex. Proc. Natl. Acad. Sci. U.S.A. 95, 15741–15746 (1998).

50. Voigts, J. & Harnett, M. T. Somatic and Dendritic Encoding of Spatial Variables in Retrosplenial Cortex Differs during 2D Navigation. Neuron 105, 237-245.e4 (2020).

51. Wullimann, M. F., Rupp, B. & Reichert, H. Neuroanatomy of the Zebrafish Brain. (Birkhäuser Basel, Basel, 1996). doi:10.1007/978-3-0348-8979-7.

52. Janiak, F. K. et al. Non-telecentric two-photon microscopy for 3D random access mesoscale imaging. Nat Commun 13, 544 (2022).

53. Schuetzenberger, A. & Borst, A. Seeing Natural Images through the Eye of a Fly with Remote Focusing Two-Photon Microscopy. iScience 23, 101170 (2020).

54. Pologruto, T. A., Sabatini, B. L. & Svoboda, K. ScanImage: Flexible software for operating laser scanning microscopes. BioMed Eng OnLine 2, 13 (2003).

55. Prevedel, R. et al. Fast volumetric calcium imaging across multiple cortical layers using sculpted light. Nat Methods 13, 1021–1028 (2016).

56. Rupprecht, P. et al. Centripetal integration of past events in hippocampal astrocytes regulated by locus coeruleus. Nat Neurosci 27, 927–939 (2024).

57. Duss, S. N. et al. Hippocampal neuronal and astrocytic responses to noradrenaline and natural arousal, eLife 15:RP111593. Reviewed preprint at 10.7554/eLife.111593.1 (2026).

58. Bethge, P. et al. An R-CaMP1.07 reporter mouse for cell-type-specific expression of a sensitive red fluorescent calcium indicator. PLOS ONE 12, e0179460 (2017).

59. Harrison, K. Optiland. Zenodo 10.5281/ZENODO.14588961 (2026).

60. Pnevmatikakis, E. A. & Giovannucci, A. NoRMCorre: An online algorithm for piecewise rigid motion correction of calcium imaging data. Journal of Neuroscience Methods 291, 83–94 (2017).

61. Rupprecht, P. & Friedrich, R. W. Precise Synaptic Balance in the Zebrafish Homolog of Olfactory Cortex. Neuron 100, 669-683.e5 (2018).

62. Li, X. et al. Reinforcing neuron extraction and spike inference in calcium imaging using deep self-supervised denoising. Nat Methods 18, 1395–1400 (2021).

63. Li, X. et al. Real-time denoising enables high-sensitivity fluorescence time-lapse imaging beyond the shot-noise limit. Nat Biotechnol 41, 282– 292 (2023).

64. Lein, E. S. et al. Genome-wide atlas of gene expression in the adult mouse brain. Nature 445, 168–176 (2007).

